# Deep learning to decipher the progression and morphology of axonal degeneration

**DOI:** 10.1101/2020.08.26.269092

**Authors:** Alex Palumbo, Philipp Grüning, Svenja Kim Landt, Lara Eleen Heckmann, Luisa Bartram, Alessa Pabst, Charlotte Flory, Maulana Ikhsan, Sören Pietsch, Reinhard Schulz, Christopher Kren, Norbert Koop, Johannes Boltze, Amir Madany Mamlouk, Marietta Zille

**Affiliations:** Fraunhofer Research and Development Center for Marine and Cellular Biotechnology, Fraunhofer Research Institution for Individualized and Cell-Based Medical Engineering, Lübeck, Germany; Institute for Medical and Marine Biotechnology, University of Lübeck, Lübeck, Germany; Institute for Experimental and Clinical Pharmacology and Toxicology, University of Lübeck, Lübeck, Germany; Institute for Neuro- and Bioinformatics, University of Lübeck, Lübeck, Germany; Department of Neonatology, Universitätsklinikum Leipzig, Leipzig, Germany; Wissenschaftliche Werkstätten, University of Lübeck, Lübeck, Germany; Medical Laser Center Lübeck GmbH, Lübeck, Germany; School of Life Sciences, The University of Warwick, Gibbet Hill Campus, Coventry, United Kingdom

**Keywords:** axon, brain hemorrhage, cell culture, machine learning, microfluidic, microscopy, stroke, time-lapse

## Abstract

**Background:** Axonal degeneration (AxD) is a pathological hallmark of many neurodegenerative diseases. Deciphering the morphological patterns of AxD will help to understand the underlying mechanisms and to develop effective therapeutic interventions. Here, we evaluated the progression of AxD in cortical neurons using a novel microfluidic device in combination with a deep learning tool, the EntireAxon, that we developed for the enhanced-throughput analysis of AxD on microscopic images.

**Results:** The EntireAxon convolutional neural network sensitively and specifically segmented the features of AxD, including axons, axonal swellings, and axonal fragments, and its performance exceeded that of human expert raters. In an *in vitro* model of AxD in hemorrhagic stroke induced by the hemolysis product hemin, we detected the concentration- and time-dependent degeneration of axons leading to a decrease in axon area, while the axonal swelling and axonal fragment area increased. Time course analysis revealed that axonal swellings preceded axon fragmentation, suggesting that swellings may be reliable predictors of AxD. Using a recurrent neural network, we further identified four morphological patterns of AxD (granular, retraction, swelling, and transport degeneration) in cortical axons subjected to hemin.

**Conclusions:** These findings indicate a morphological heterogeneity of AxD under pathophysiologic conditions. The combination of the microfluidic device with the EntireAxon deep learning tool enable the systematic analysis of AxD but also unravel a so far unknown intricacy in which AxD can occur in a disease context.

## Introduction

Axonal degeneration (AxD) is a process in which axons disintegrate physiologically during nervous system development and aging, or as a pathological element of degenerative nervous system diseases (Luo and O’Leary, 2005; Lingor et al., 2012; Salvadores et al., 2017). Apart from axonal fragments, axon swellings (also called axonal beadings, bubblings or spheroids) are a hallmark of degenerating axons (Saxena and Caroni, 2007; Lingor et al., 2012; Wang et al., 2012), containing disorganized cytoskeleton and organelles resulting from an interruption of axonal transport (Coleman, 2005; Nikić et al., 2011; Yong et al., 2019).

It is known that axons disintegrate in different ways depending on the biological context. During development and neural circuit assembly, inappropriately grown axons can undergo axonal retraction, axonal shedding or local AxD (Pease and Segal, 2014; Neukomm and Freeman, 2014). Axonal retraction is characterized by retraction bulb formation at the distal tip, and subsequent pullback (Pease and Segal, 2014). During axonal shedding, the axon retracts leaving behind small pieces of its distal part (axosomes) (Bishop et al., 2004). Local AxD is characterized by axon disintegration into separated axonal fragments (Neukomm and Freeman, 2014). Acutely and chronically injured axons may degenerate retrogradely (distal-to-proximal direction, dying-back), anterogradely (proximal-to-distal direction) or in a Wallerian degeneration pattern (distal part of the axon from injury site), ultimately resulting in the generation of axonal fragments (Cavanagh, 1979; Coleman, 2005; Beirowski et al., 2005). However, AxD patterns have been mainly described in extracerebral axons in models of nutrient deprivation or axotomy.

Not much is known on AxD in cortical neurons subjected to a disease-specific cytotoxic micromilieu. A distinct pathological micromilieu has recently been observed for hemorrhagic stroke, after which the lysis of erythrocytes from the hematoma leads to the release of the cytotoxic product hemin (Robinson et al., 2009; Zille et al., 2017). Patients suffering from hemorrhagic stroke often experience AxD that is associated with worse motor and functional outcome (Venkatasubramanian et al., 2013; Chen et al., 2018). Importantly, AxD occurs in the subacute stages of hemorrhagic stroke. Thus, addressing AxD may not only provide a new therapeutic target, but also a much wider time window for intervention. Since not much is known about the mechanisms, morphological patterns, and the temporal progression of AxD in the context of hemorrhagic stroke, we here sought to examine the progression of AxD and its associated morphological alterations.

As the disintegration of the axons endures from minutes to hours (Beirowski et al., 2005; Kerschensteiner et al., 2005), it is necessary to monitor the spatiotemporal progression of AxD and its morphological hallmarks continuously. However, conventional software solutions fail to automatically detect and quantify high axon numbers as well as axonal swellings and fragments in phase-contrast microscopic images. The reason may be two-fold: 1) Conventional software relies on image binarization (Sasaki et al., 2009; Becker and Madany, 2012), which can lead to information loss and low sensitivity as thin axons may not be recognized. 2) The analysis requires subjective and time-consuming manual annotations, e.g., thresholding and defining the region of interest (Pool et al., 2008; Ho et al., 2011; Li et al., 2014). So far, immunostained images were used to investigate morphological changes in AxD as the analysis of phase-contrast images has been limited by the lower target-to-background signal. Immunofluorescence images, however, entail certain disadvantages such as photobleaching and the requirement for cell fixation, which restricts observations to a single time point. Thus, a software tool for the automatized detection and quantification of the morphological patterns of AxD in long-term live cell imaging is required to improve both sensitivity and throughput to overcome current limitations in understanding AxD. In this study, we demonstrate that cortical axons underwent AxD after the exposure to the hemolysis product hemin, with axonal swellings preceding axon fragmentation. Deep learning further detected the occurrence of four AxD patterns being characterized as granular, retraction, swelling, and transport degeneration. This may inform downstream AxD and neurodegeneration research in health and disease. We also provide tools for the enhanced throughput analysis of AxD, including a microfluidic device containing 16 independent experimental units and the deep learning platform “EntireAxon” to analyze AxD, which will help augment our understanding of AxD and may also support the development of novel treatment approaches for neurodegenerative diseases.

## Results

### An enhanced throughput microfluidic device and the EntireAxon deep learning tool allow the longitudinal study of axonal degeneration

The major limiting factor of commercially available microfluidic devices to study AxD is that they are single, individual systems and hence, can only be used to assess one condition, which is time-consuming and precludes high-throughput analyses. To enable the systematic analysis of AxD *in vitro*, we 1) manufactured a microfluidic device containing 16 individual microfluidic units (**Fig. 1 and Supplementary Fig. S1**) that can be investigated in parallel and recorded simultaneously, and 2) trained a convolutional neural network (CNN), the EntireAxon, to segment all relevant features of AxD, i.e., axons, axonal swellings, and axonal fragments (**Fig. 2**).

**Figure 1.**
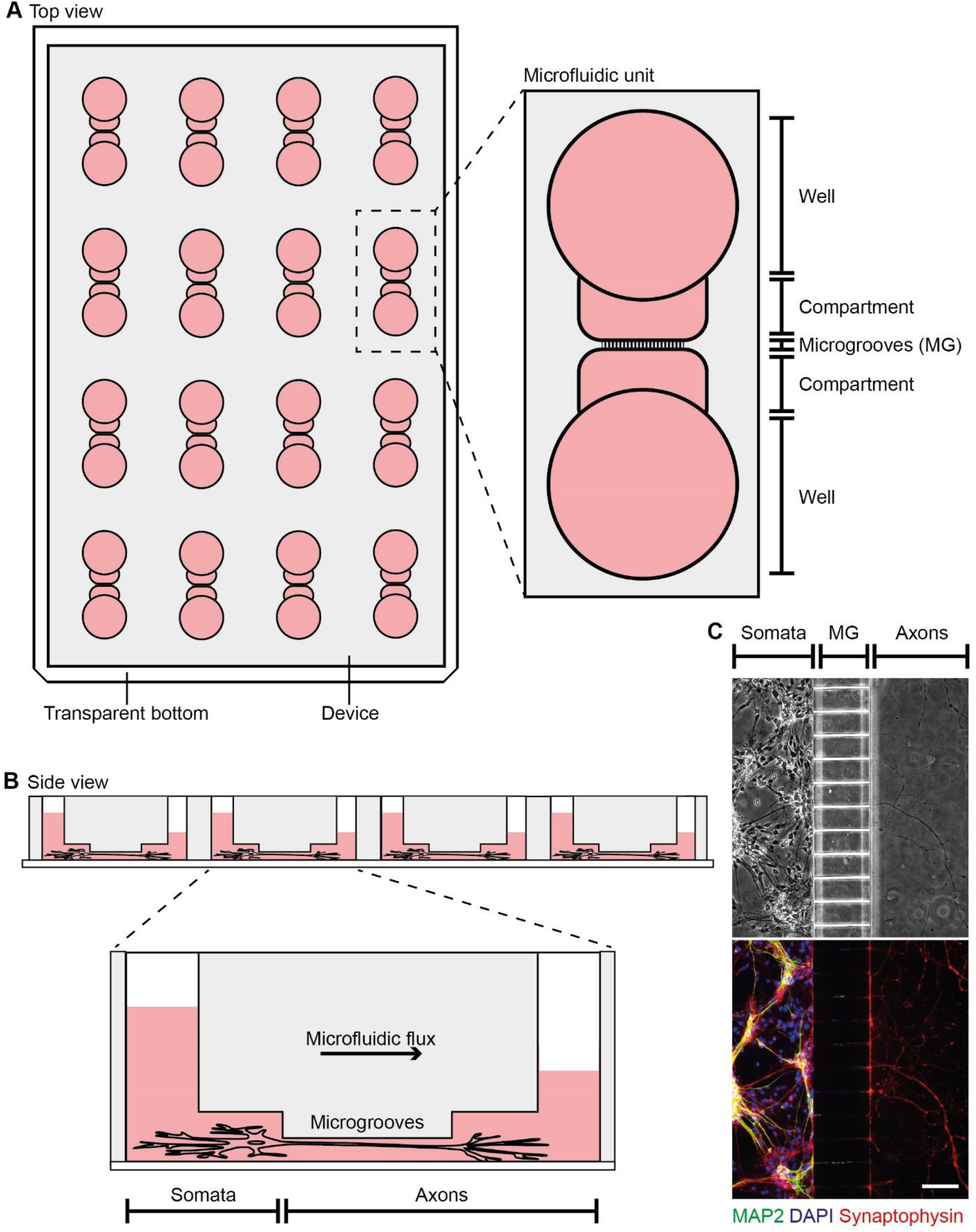
Microfluidic device for the enhanced throughput cultivation of axons. **(A)** The microfluidic device incorporates 16 individual microfluidic units for axon cultivation. One microfluidic unit consists of two wells that are connected through compartments and microgrooves (MG). **(B)** Primary cortical neurons are seeded into the soma compartment from which their axons grow through the MG into the axon compartment. Directed growth is supported by culture medium microflux due to different medium volumes between the two wells. **(C)** Phase-contrast image of primary cortical axons that were spatially separated from their somata by the MG at day *in vitro* 7, which we confirmed by immunofluorescence staining of dendrites using microtubule-associated protein 2 (MAP2, green, 1:4000) and axons using synaptophysin (red, 1:250). DAPI (blue, 1:1000)) was used for nuclear counterstaining (top). Scale bar: 100 μm.

**Figure 2.**
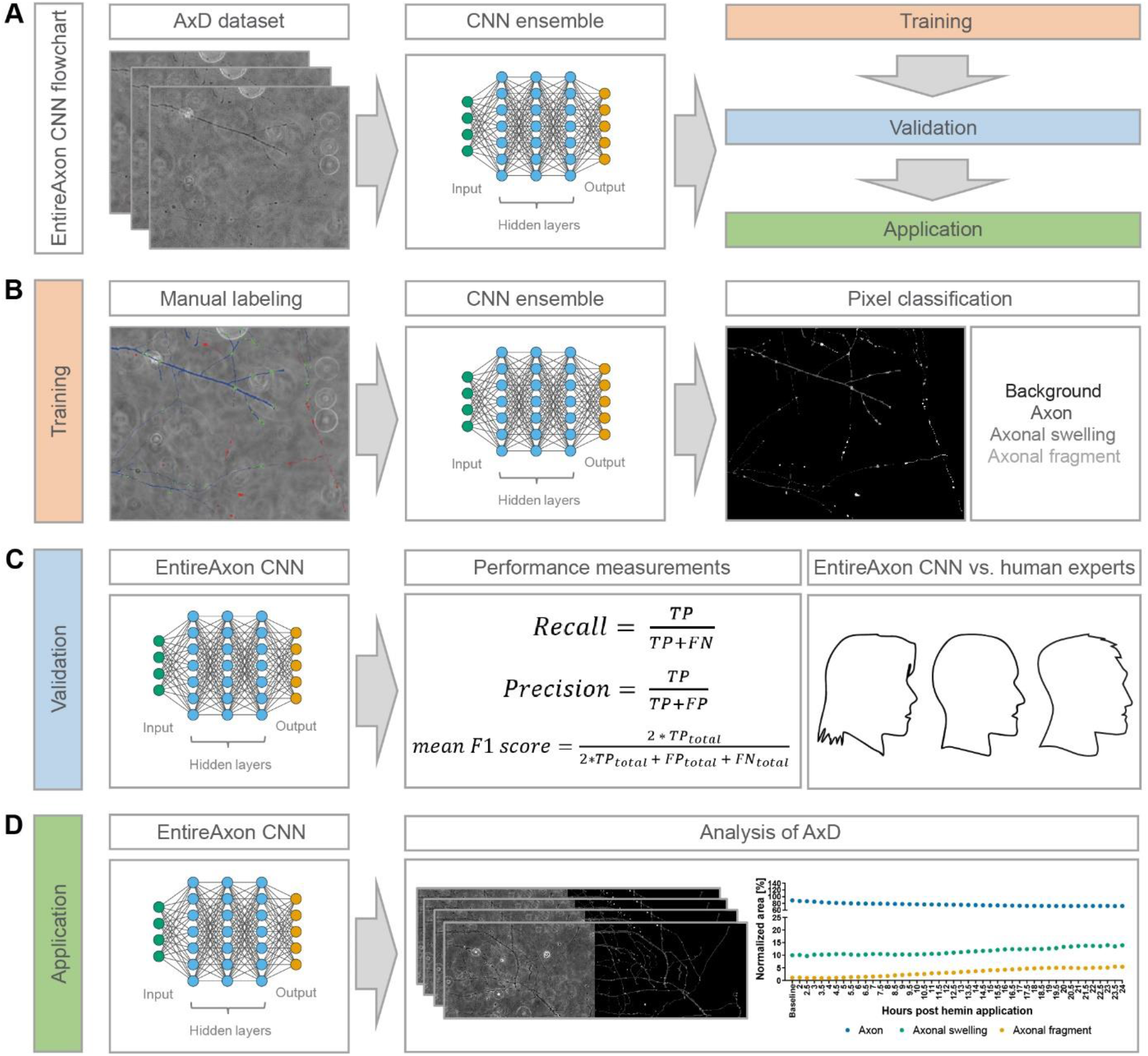
EntireAxon CNN for the enhanced throughput analysis of AxD. **(A)** The flow chart of the EntireAxon CNN. The AxD data was separated into training, validation, and testing data. We adapted a standard u-net with ResNet-50 encoder (Ronneberger et al., 2015; He et al., 2015) and used a CNN ensemble, which combines predictions from multiple CNNs to generate a final output and is superior to individual CNNs (Dietterich, 2000; Huang et al., 2016; Vuola et al., 2019). **(B)** We manually labeled the training data to segment each pixel into the four classes ‘background’, ‘axon’, ‘axonal swelling’, and ‘axonal fragment’, which are displayed in the output image in black, dark grey, intermediate grey, and light grey, respectively. We trained an ensemble comprising 8 CNNs to segment the four classes. **(C)** The EntireAxon CNN was validated with a separate validation dataset to assess its performance (recall, precision, and mean F1 score), which was compared to human experts (ground truth was labeled by human expert 1). **(D)** The EntireAxon CNN was applied to data on AxD induced by the exposure of hemin, which is used to model of hemorrhagic stroke *in vitro*.

While the EntireAxon CNN recognized the class ‘background’ better than the three axon classes ‘axon’, ‘axonal swelling’, and ‘axonal fragment’ (mean F1 score: 0.995), axon-specific segmentation revealed the highest mean F1 score for the class ‘axon’ (0.780), followed by the classes ‘axonal swelling’ (0.567), and ‘axonal fragment’ (0.301) (**Fig. 3A**). Next, we compared the performance of the EntireAxon CNN on the ground truth (human expert 1) with two additional human experts. The EntireAxon CNN reached higher mean F1 scores for all classes, except for the class ‘axonal fragment’, where human expert 2 outperformed the EntireAxon CNN (**Fig. 3B**). This may have been due to the fact that the EntireAxon CNN was trained on images labeled by the same human expert (1) that labeled the ground truth. To assess whether its performance is more generalizable across the different experts, we compared the EntireAxon CNN to each of the human experts on the consensus labels of the two other human experts (**Fig. 3C-D**). Visual inspection of the labels showed a wide overlap between the different experts, but also that there was considerable uncertainty, especially for the classification of axonal fragments (**Fig. 3C**). When comparing the mean F1 scores for all classes, the EntireAxon reached similar or even higher scores than the other three experts (**Fig. 3D**). Collectively, this suggests that the EntireAxon CNN sensitively and specifically recognizes axons and the morphological features of AxD.

**Figure 3.**
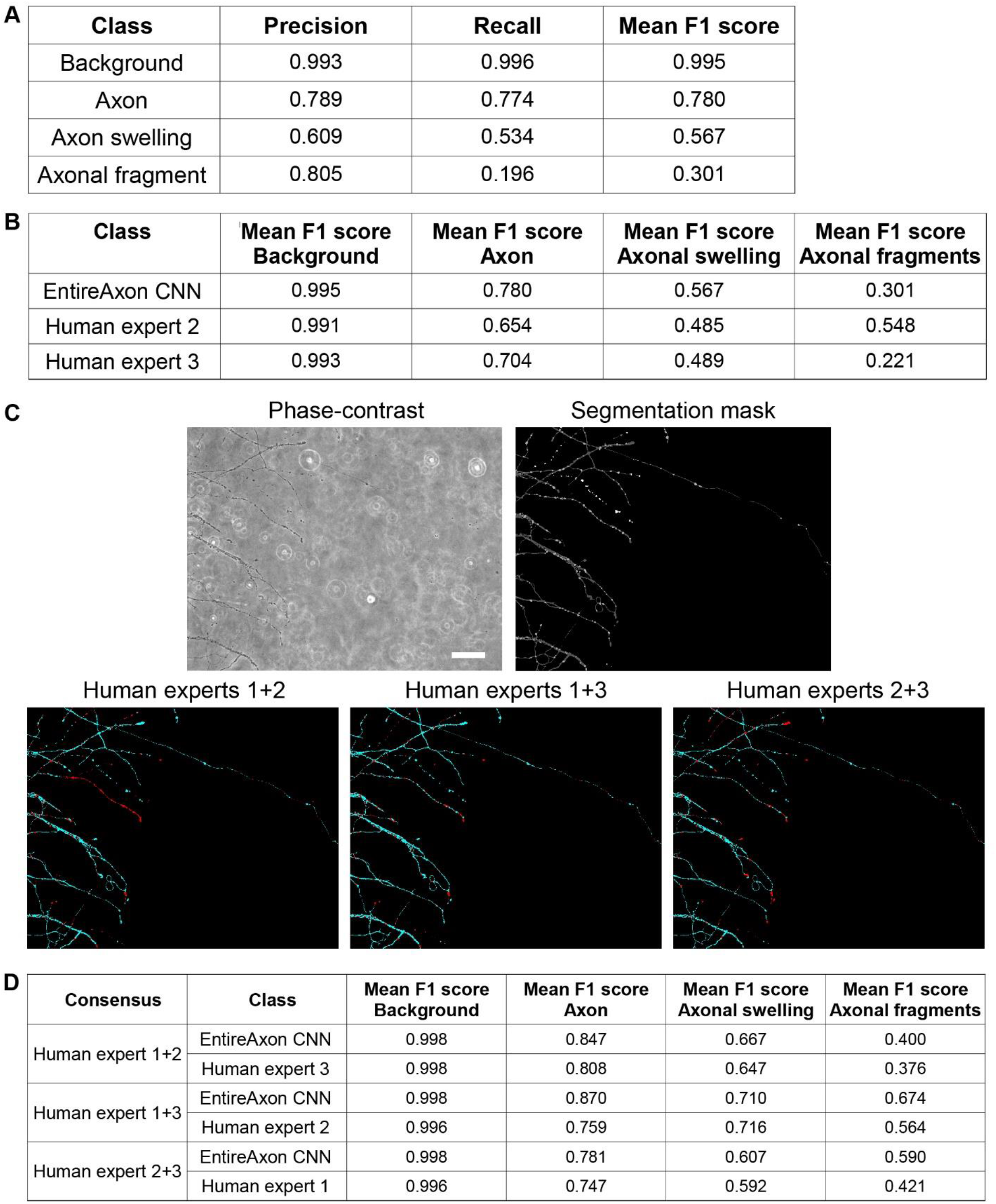
Performance of the EntireAxon CNN compared to human experts. **(A)** Validation of the EntireAxon CNN performance for all four classes ‘background’, ‘axon’, ‘axonal swelling’ and ‘axonal fragment’ in before unseen phase-contrast microscopic images. **(B)** Comparison of the mean F1 scores between the EntireAxon CNN and two human experts on the ground truth (human expert 1 who also labeled the training images) to recognize background, axon, axonal swelling and axonal fragments. **(C)** Phase-contrast validation image, its EntireAxon CNN segmentation mask, and the consensus labeling masks of two human experts that show the segmentation overlap (cyan) or difference (red) between the labels. Scale bar: 100 μm. **(D)** Comparison of the mean F1 scores between the EntireAxon CNN and the human expert on the consensus labeling of the other two human experts.

### Axonal integrity is lost over time with axonal swellings preceding axon fragmentation

We then applied the EntireAxon CNN to assess AxD in the context of hemorrhagic stroke. We applied the hemolysis product hemin, a commonly used agent to mimic hemorrhagic stroke *in vitro* (Robinson et al., 2009; Zille et al., 2017; Chen and Regan, 2004), on primary cortical axons. Accordingly, isolated axons were exposed to the hemolysis product hemin and recorded by time-lapse microscopy for 24 hours. Hemin induced concentration- and time-dependent morphological changes leading to AxD compared to vehicle-treated axons (**Fig. 4 and Videos S1-4**). Area under the curve (AUC) analyses revealed a significant decrease in axon area in all three hemin concentrations (50 μM vs. 0 μM: *P* = 0.026; 100 μM vs. 0 μM: *P* = 0.018, 200 μM vs. 0 μM: *P* < 0.001). The axonal swelling area also increased in all three concentrations (50 μM vs. 0 μM: *P* = 0.012, 100 μM vs. 0 μM: *P* = 0.005, 200 μM vs. 0 μM: *P* = 0.016), while the axonal fragment area was elevated only for axons treated with 100 and 200 μM hemin (vs. 0 μM: *P* = 0.004, **Fig. 5 and Table S2**).

**Figure 4.**
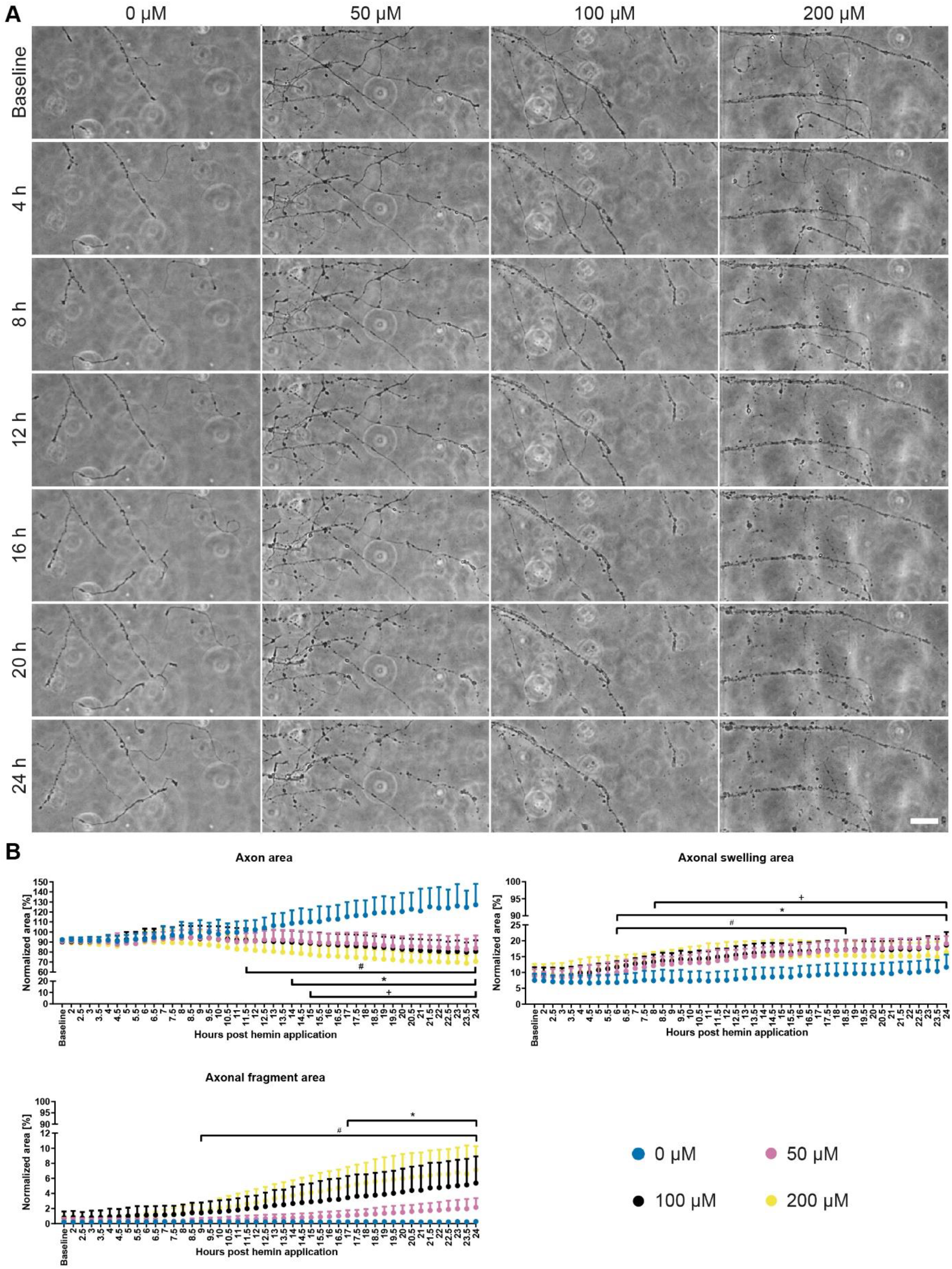
Time- and concentration-dependent hemin-induced AxD. **(A)** Primary cortical axons treated with hemin (50, 100, 200 μM) degenerated compared to vehicle-treated axons (0 μM) that continued to grow. Scale bar: 50 μm. For complete time-lapse videos including segmentation, refer to **Video S1-4**. **(B)** Quantification of AxD over 24 hours in phase-contrast images. To determine the time course, the sum of pixels in each class and hemin concentration over time was normalized to the baseline of that class and condition. The quantification of the phase-contrast images over 24 hours revealed significantly smaller axon areas starting at 11.5 hours after 200 μM (*P* = 0.020), at 14 hours after 100 μM (*P* = 0.040), and at 15 hours after 50 μM (*P* = 0.018) hemin treatment compared to control (0 μM). The axonal fragment area significantly increased from 9.5 hours onwards in 200 μM hemin (*P* = 0.037) and from 17.5 hours in 100 μM hemin (*P* = 0.044), while the axonal swelling area increased from 6 hours onwards in 100 μM hemin (*P* = 0.019) and 200 μM hemin (*P* = 0.010) and from 8 hours in 50 μM hemin (*P* = 0.030). N = 6 independent cultures of primary cortical neurons. Means + 95 % CI are given. One-way ANOVA with Greenhouse-Geisser correction. +, *, # *P* < 0.05; + = 50 μM vs. 0 μM, * = 100 μM vs. 0 μM, # = 200 μM vs. 0 μM. For detailed statistical information, refer to **Table S1**.

**Figure 5.**
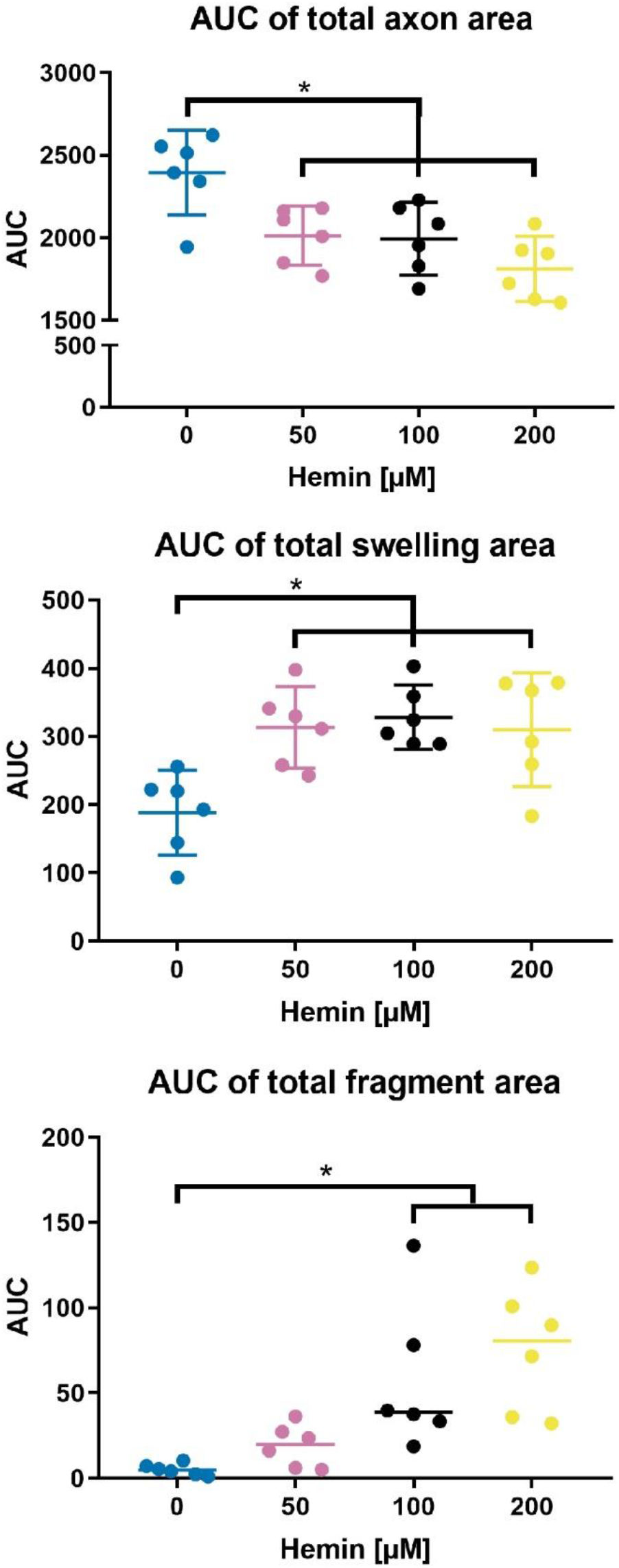
Area under the curve (AUC) analysis of hemin-induced AxD. While axons exposed to hemin showed a decline in axon area, axonal swelling and axonal fragment area increased. N = 6 independent cultures of primary cortical neurons. Means ± 95 % CI are given for axon and axonal swelling area, medians for fragment area. * *P* < 0.05 vs. 0 μM for axon and swelling area, * *P* < 0.0167 for fragment area due to manual Bonferroni correction for nonparametric data. For exact *p* values, refer to **Table S2**.

Comparing the time course of AxD between hemin- and vehicle-treated axons (0 μM), the axon area decreased starting at 11.5 hours at 200 μM (*P* = 0.020, from 15 hours *P* < 0.001), at 14 hours at 100 μM (*P* = 0.040, from 18.5 hours *P*<0.001), and at 15 hours at 50 μM (*P* = 0.018, from 19 hours *P* < 0.001). Hemin treatment also elevated the axonal fragment area starting at 9 hours at 200 μM (*P* = 0.037) and at 17 hours at 100 μM hemin (*P* = 0.044). Interestingly, the axonal swelling area increased prior to the changes in axon and axonal fragment area, i.e., starting at 6 hours at 200 μM (*P* = 0.010) and 100 μM (*P* = 0.019), and at 8 hours at 50 μM hemin (*P* = 0.030). For the highest hemin concentration, the increase was only transient (until 18.5 hours), suggesting that axonal swellings preceded the axon fragmentation (**Table S1**), which can also be seen in the time-lapse recordings (**Videos S2-4**).

The results of the time course analysis were further substantiated by live cell fluorescent staining (calcein AM), which indicated the starting point of AxD after hemin treatment between 8 and 12 hours for 200 μM hemin, between 12 and 16 hours for 100 μM hemin and 16 and 20 hours for 50 μM hemin (**Supplementary Fig. S2**). Taken together, AxD progression depends on the severity of the insult and axonal swellings may be reliable predictors of AxD.

### Deep learning deciphers four patterns of AxD

AxD time-lapse data revealed different morphological patterns of degeneration that can occur in the same axons over time (**Fig. 6** and **Videos S5-8**). We categorized these morphological patterns as:

i. Granular degeneration: AxD resulting in granular separated fragments.
ii. Retraction degeneration: AxD in which the distal part of the axon retracts ultimately resulting in granular degeneration.
iii. Swelling degeneration: AxD in which axonal swellings enlarge, followed by granular degeneration.
iv. Transport degeneration: AxD in which axonal swellings of constant size, which do not enlarge, are transported along the axon resulting in granular degeneration.

**Figure 6.**
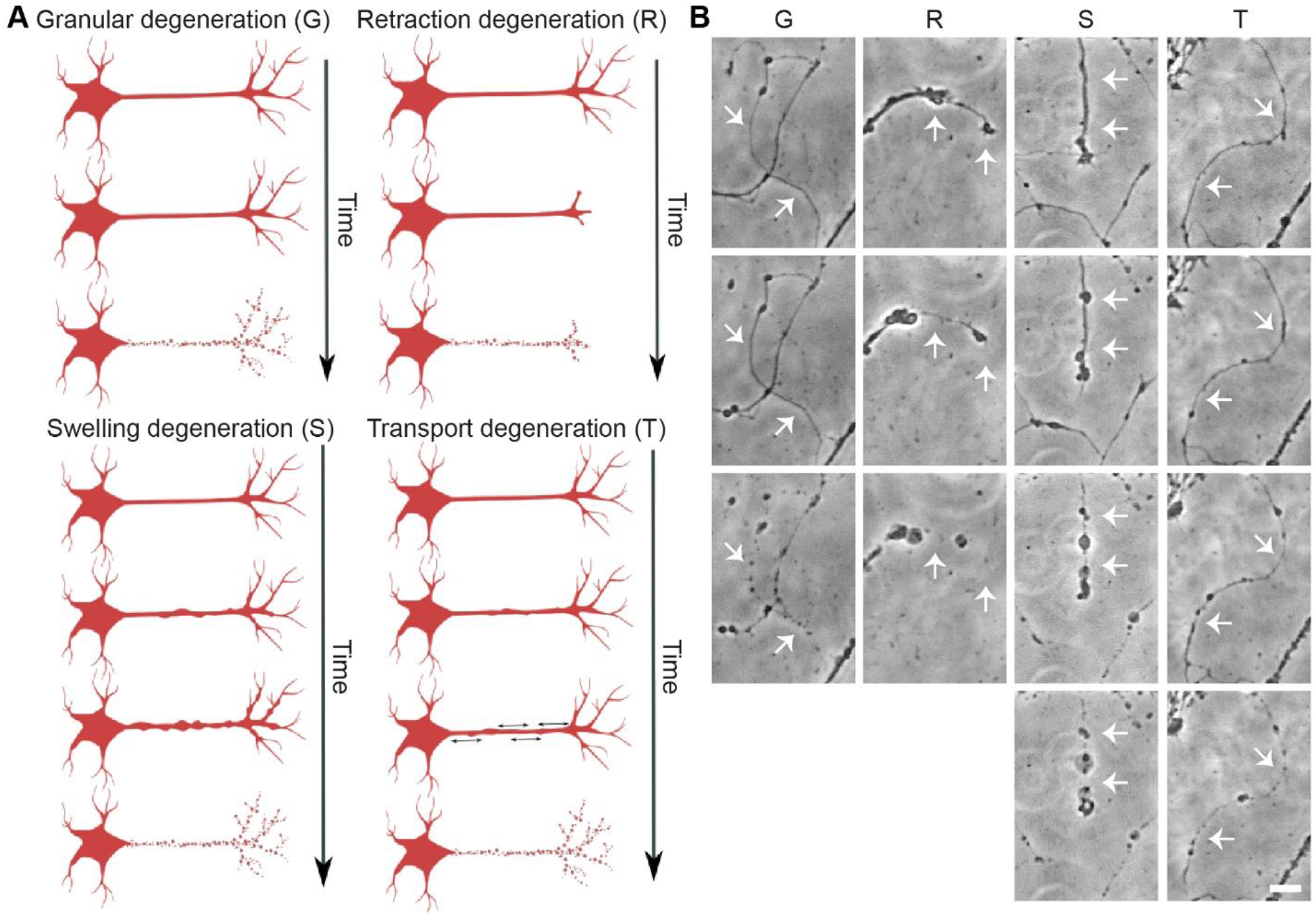
Four morphological patterns of AxD. **(A)** Schematic overview of the proposed AxD morphological patterns: granular degeneration, retraction degeneration, swelling degeneration, and transport degeneration. **(B)** Phase-contrast recordings of the four morphological patterns of AxD in primary cortical axons. Granular degeneration (G) is characterized by the fragmentation of the axon (white arrows). During retraction degeneration (R), the axonal growth cone retracts in the proximal direction and the part of the axon in proximity of the growth cone disintegrates accompanied by axonal swellings (white arrows). During swelling degeneration (S), many axonal swellings enlarge resulting in axonal fragments (white arrows). During transport degeneration (T), axonal swellings are transported along the axon prior to the degeneration of the axon (white arrows). Scale bar: 20 μm. For complete time-lapse videos including segmentation, refer to **Videos S5-8.**

We trained a recurrent neural network (RNN), the EntireAxon RNN, to identify these morphological patterns based on changes in class segregation over time using a training dataset of AxD segmentation recordings (**Fig. 7A**). Given the four different classes (background, axon, axonal swelling, and axonal fragment), 16 different class pairs can occur between a segmentation at time step *t* and time step *t*+1. For example, a background pixel at *t* can either remain background pixel at *t*+1 or change into one of the other three classes, and the same is true for the other classes. Thus, in total, four times four class pairs are possible. We used a window size of 32×32, of which always the probability of a class pair in the central pixel relative to the previous time point was computed for each time point and across the entire image.

**Figure 7.**
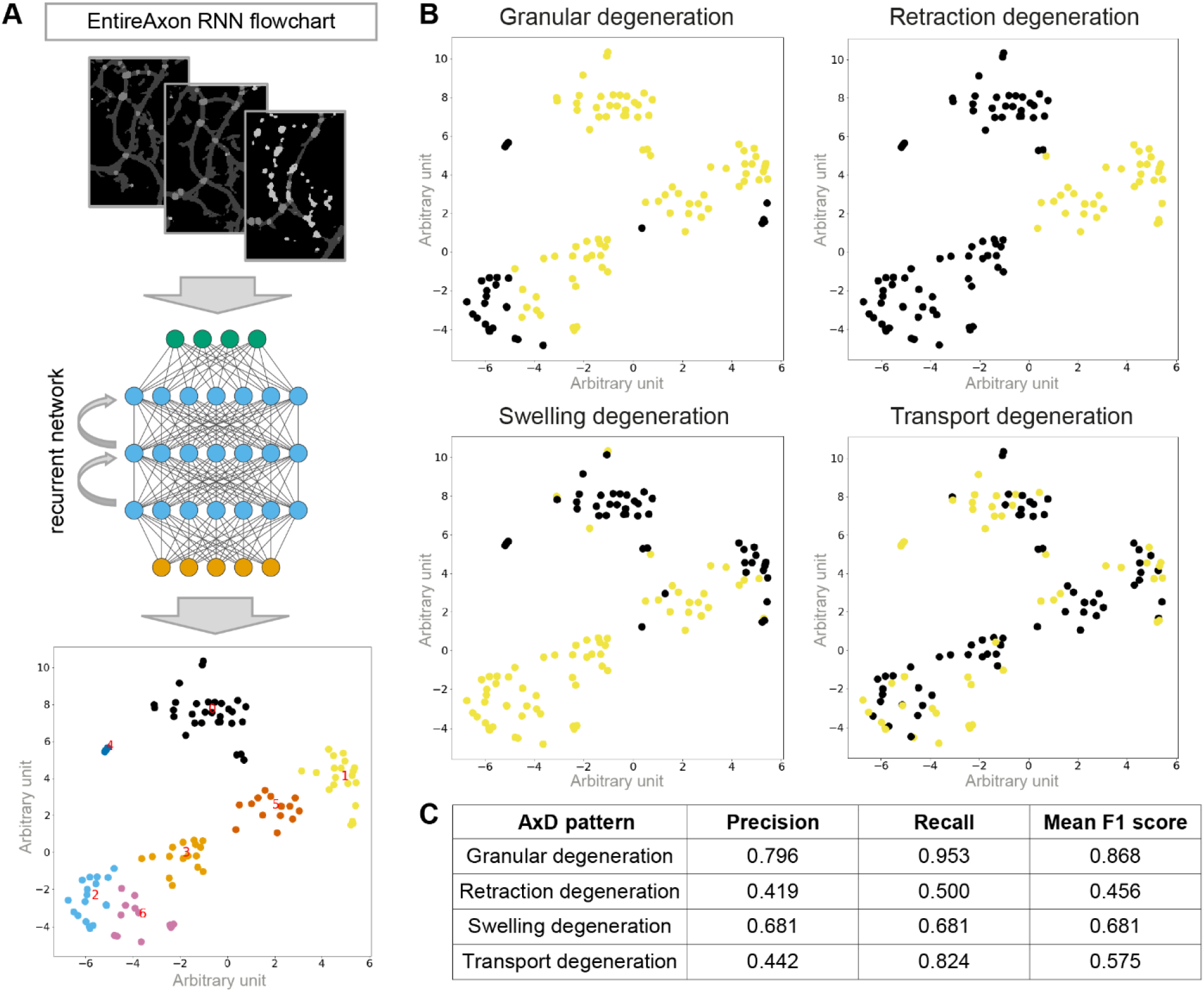
Recognition of four morphological patterns of AxD by the EntireAxon RNN. **(A)** Schematic workflow of the RNN to recognize and quantify morphological patterns of AxD based on the identification of seven clusters. The EntireAxon CNN segmentation masks were used for the RNN training, which determined the change in class over time. Based on the 16 different possible class pairs, the RNN determined seven clusters (cluster 0-6). To visualize the relationships of the specific samples, we employed t-distributed stochastic neighborhood embedding (T-SNE) to compute a 2-dimensional representation of the high-dimensional data. **(B)** The clusters classify the four morphological patterns of AxD with yellow indicating included and purple indicating excluded clusters: granular (G), retraction (R), swelling (S), and transport degeneration (T). Clusters of granular degeneration overlap with recognized clusters of other morphological patterns (retraction, swelling, and transport degeneration). For more details on the morphological changes underlying the cluster analysis, refer to **Supplementary Fig. S3**. **(C)** 10-fold cross-validation of the four morphological patterns of AxD.

The RNN determined seven clusters (cluster 0-6) that were characterized by an idiosyncratic pattern of changes in class distribution over 24 hours (**Supplementary Fig. S3**). All clusters showed a decrease in the class ‘axon’ and an increase in the class ‘background’. Depending on the hemin concentration, the changes occurred at a different magnitude and at different time points, and with concomitant increases in either the class ‘axonal swelling’ and/or ‘axonal fragment’. In cluster 0, there was an early decrease in the class ‘axon’, which then continued more linearly as well as a later rise in the class ‘axonal fragment’. In contrast to cluster 0, cluster 1 showed no increase in the class ‘axonal fragment’ and a linear decrease in the class ‘axon’ from the start. In cluster 2, there was a strong increase in the class ‘axonal swelling’. Cluster 3 demonstrated an early and lasting high level of the class ‘axonal swelling’ with a later increase in the class ‘axonal fragment’. Cluster 4 showed a rapid decrease in the class ‘axon’ concomitant with an increase in the classes ‘background’ and ‘ axonal swelling’. Cluster 5 was similar to cluster 1, but with an early drop in the class ‘axon’. Cluster 6 showed an increase in the class ‘axonal swelling’ similar to but to a greater extent than cluster 2.

The RNN categorized each cluster to one of the four morphological patterns (**Fig. 7B**): i) Granular degeneration was defined by clusters that describe the degeneration of axons into axonal fragments, i.e. clusters 0, 1, 3, and 5. ii) Retraction degeneration only included the clusters 1 and 5, indicating the retraction of the axon followed by its fragmentation. iii) Swelling degeneration was characterized by the three clusters that included the class ‘axonal swelling, i.e., clusters 2, 3, and 6, as well as cluster 5 showing the exchange of the class ‘axon’ for ‘background’. iv) Transport degeneration was the only pattern that relied on cluster 4 and was also characterized partly on clusters 0, 1, 2, and 6. Although some clusters overlap among morphological patterns, the unique combination of the different clusters allows to distinguish all four morphological patterns.

To validate the EntireAxon RNN, a 10-fold cross-validation was performed. Therefore, the dataset was randomly divided into 10 datasets and ten models were trained with 9 of the datasets leaving the remaining dataset for validation (not previously seen by the RNN). Based on the combined test samples, the RNN was able to distinguish between the four morphological patterns of AxD (**Fig. 7C**). These data confirm that the combination of the different AxD features as well as their spatiotemporal progression defines distinct morphological AxD patterns.

### The morphological patterns of AxD depend on the extent of AxD

We then applied the EntireAxon RNN to quantify the occurrence of the four morphological patterns of AxD in the context of hemorrhagic stroke (**Fig. 8 and Video S9**). While all AxD patterns were detected (**Fig. 8A**), hemin concentration-dependently increased granular degeneration (*P* < 0.001), swelling degeneration (*P* < 0.001), and transport degeneration (*P* = 0.025, **Fig. 8B**). When comparing the slopes of the different AxD patterns under hemin exposure, granular and swelling degeneration were significantly different from transport degeneration (*P* = 0.005 and *P* = 0.004, respectively, **Table S3**). Collectively, our data suggest that hemin concentration-dependently induces different morphological patterns of AxD in cortical axons.

**Figure 8.**
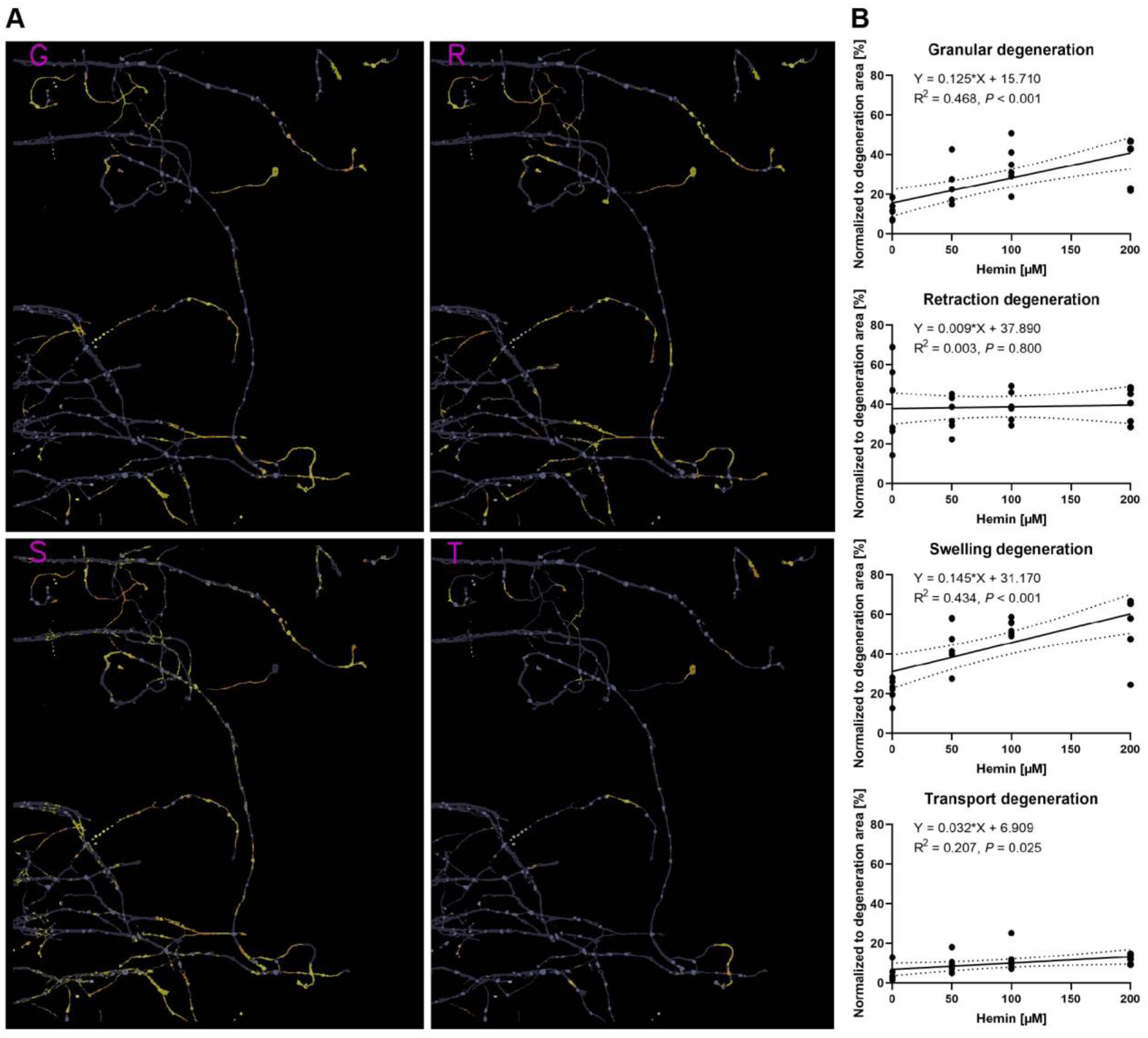
Concentration-dependent differences in the morphological patterns of hemin-induced AxD. **(A)** The classification of granular (G), retraction (R), swelling (S), and transport degeneration (T) in primary cortical axons treated with 200 μM hemin. For the complete time-lapse video including segmentation, refer to **Video S9. (B)** Linear regressions of the four morphological patterns of AxD in hemin-induced AxD. The area classified for each AxD pattern was normalized to the total degeneration area. Dotted lines show 95 % confidence bands. N = 6 independent cultures of primary cortical neurons. Granular degeneration: F(1,22) = 19.330, *P* < 0.001. Retraction degeneration: F(1,22) = 0.066, *P* = 0.800. Swelling degeneration: F(1,22) = 16.900, *P* < 0.001. Transport degeneration: F(1,22) = 5.757, *P* = 0.025. For the comparison of the slopes between the different AxD patterns, refer to **Table S3**.

## Discussion

We here describe the occurrence of four morphological patterns of AxD under pathophysiological conditions: granular, retraction, swelling, and transport degeneration. These rely on time- and concentration-dependent changes of the morphological features of AxD, with axonal swellings preceding axon fragmentation. The herein introduced complementary tools, a novel microfluidic device and the EntireAxon, allow increasing the experimental yield, the in-depth enhanced throughput analysis of AxD as well as the longitudinal investigation of AxD.

We propose a novel monolithic microfluidic device consisting of 16 individual microfluidic units that enables the parallel and separated treatment and/or manipulation of axons and somata (**Fig. 1**). The currently available devices do not allow enhanced throughput experiments as they comprise only single microfluidic units (Park et al., 2006; Van Laar et al., 2019). Although some devices can harbor multiple experimental conditions, they employ a radial design with a single soma compartment, in which one experimental condition may influence another due to the potential of retrograde signaling (Hosmane et al., 2010; Biffi, 2015). Another option is the parallel use of multiple individual devices, which allows handling up to 12 devices in a conventional 12-well plate (Li et al., 2014). Compared to our device, this procedure is time-consuming in both the manufacturing and adjustment for recordings.

The extent of AxD has so far been mainly investigated with a focus on axon fragmentation as primary readout. To quantify axon fragmentation, Sasaki and colleagues introduced the AxD index as the ratio of fragmented axon area versus total axonal area (Sasaki et al., 2009). However, the AxD index did not include axonal swellings, which are a characteristic feature of degenerating axons (Yong et al., 2019; Cui et al., 2020). Although other analyses considered axonal swellings as a morphological feature of AxD (Nikić et al., 2011; Yong et al., 2019), the approaches were time-consuming and required manual annotations.

We herein adapted a standard u-net with ResNet-50 encoder (Ronneberger et al., 2015; He et al., 2015) and used a CNN ensemble, which combines predictions from multiple CNNs to generate a final output and is superior to individual CNNs (Dietterich, 2000; Huang et al., 2016; Vuola et al., 2019). The EntireAxon CNN performs an automatic segmentation and quantification of axons and morphological features relevant to AxD, including axonal swellings and fragments, on phase-contrast time-lapse microscopy images (**Fig. 2**). The EntireAxon CNN recognized the four classes ‘background’, ‘axon’, ‘axonal swelling’, and ‘axonal fragment’, with the highest mean F1 score for the class ‘background’ (**Fig. 3A**). The comparably lower performance of the CNN to recognize axonal fragments may be explained by the disproportional distribution of pixels in the training and validation data (‘background’ mean of 96.42 % of pixels, ‘axon’ 2.77%, ‘axonal swelling’ 0.58%, ‘axonal fragment’ 0.23 %). Hence, every individual segmentation error more strongly affects the false positive or false negative rate in these classes.

Comparison with human experts revealed that the EntireAxon CNN reached a similar performance level. As expected, its performance was slightly better than the human experts on the ground truth as both, ground truth and training data, were labeled by the same human expert (**Fig. 3B**). Interestingly, when comparing the EntireAxon CNN with a human expert on the consensus label of the other two human experts, not only was the EntireAxon CNN as good as or even better than the human expert, but the mean F1 scores were also higher than on the ground truth labels (**Fig. 3D**). This may be because pixels that were differentially assigned by the human expert, i.e. more difficult to classify, were excluded from the comparison. Taken together, these findings demonstrate that the EntireAxon CNN is suitable to automatically quantify AxD and its accompanying morphological changes in an enhanced throughput manner.

Conventional *in vitro* models of AxD rely mainly on nutrient deprivation or axotomy and focus on axons outside the brain. However, AxD is not only an active and commonly observed process in the brain, but it is also believed to be caused by more complex mechanisms given the different microenvironments in which it may occur. For example, AxD has been demonstrated to occur in intracerebral hemorrhage (Venkatasubramanian et al., 2013; Tao et al., 2017). In this context, cortical axons are exposed to a cytotoxic microenvironment due to hemolysis leading to the release of blood breakdown products, whose effects on axons remain to be elucidated (Hemorrhagic Stroke Academia Industry (HEADS) Roundtable Participants, 2018). We therefore modeled hemorrhagic stroke by exposing axons from primary cortical neurons to the hemolysis product hemin and investigated the progression of AxD. Similar to previous results where 100 μM hemin were sufficient to induce significant neuronal cell death in conventional cultures of somata and axons (Zille et al., 2017), we here observed that 100 μM hemin led to a significant decrease in axon area and an increase in axonal swelling and fragment area (**Fig. 4**).

The progression of AxD undergoes a latent phase, during which the structural integrity of the axon is maintained, followed by a catastrophic phase with the rapid disintegration of the axon (Yong et al., 2019). In our model, the catastrophic phase of AxD started within 12 to 18 hours after the administration of hemin (**Fig. 4 and Supplementary Fig. S2**). Similar durations of the latent phases of AxD have been observed in other models. For instance, under circumstances of growth factor withdrawal, the transition to the catastrophic phase occurred at 12-24 hours (Nikolaev et al., 2009; Maor-Nof et al., 2016; Yong et al., 2019).

We further demonstrated that the relative axon area decreased at higher hemin concentrations, while the axonal fragment area increased. Our results are in accordance with other experimental conditions such as axotomy-mediated or paclitaxel-induced AxD, in which axonal fragments also increased (Sasaki et al., 2009; Pease-Raissi et al., 2017). As the axonal swelling area preceded the increase of axonal fragments and axon area loss, our findings are also in line with results reported in a model of experimental autoimmune encephalomyelitis indicating that axonal swelling anticipates fragmentation (Nikić et al., 2011). This suggests that axonal swelling may be a reliable predictor of AxD.

Interestingly, axonal swellings and axonal fragments were related to different morphological patterns of AxD. Specifically, we observed axons that showed signs of axonal retraction, enlarging of axonal swellings and axonal transport before degeneration (**Fig. 6**). We therefore trained the EntireAxon RNN to quantify the occurrence of four morphological patterns of AxD, i.e. granular, retraction, swelling, and transport degeneration, based on the clusters of unique changes of classes over time (**Fig. 7 and Supplementary Fig. S3**). These patterns have not been described to occur simultaneously in the same biological condition: Granular degeneration has previously been observed in retrograde, anterograde, Wallerian and local AxD after axotomy or trophic factor deprivation (Cavanagh, 1979; Coleman, 2005; Beirowski et al., 2005; Neukomm and Freeman, 2014). Retraction degeneration has been described in axonal retraction and shedding in developmental AxD (Bishop et al., 2004; Pease and Segal, 2014). Swelling degeneration was previously reported in experimental autoimmune encephalitis and growth factor deprivation (Nikić et al., 2011; Yong et al., 2019). Transport degeneration has not been reported before. However, microtubule breaks have been demonstrated in a model of axonal stretch injury. Those developed into axonal swellings resulting in axonal transport interruption with AxD as a consequence (Tang-Schomer et al., 2012).

Our data demonstrate that all four morphological degeneration patterns can occur along cortical axons (**Fig. 8**). Interestingly, we also observed a concentration-dependent effect in the context of hemorrhagic stroke. Granular, swelling, and transport degeneration were significantly increased with increasing hemin concentrations, with granular and swelling degeneration being more strongly correlated. To what extent our model of hemin-induced AxD in hemorrhagic stroke is molecularly similar to developmental or pathophysiological AxD needs to be further investigated along with the underlying molecular mechanisms of the four patterns of AxD. This could be greatly facilitated by the EntireAxon RNN that is able to automatically detect the morphological patterns in time-lapse recording due to its capacity to relate each output to previous images in the stacks by its current units.

### Limitations and outlook

i. Our microfluidic device currently does not allow to investigate AxD at more proximal axonal parts to the soma such as the axonal initial segment. Shortening the length of the microgrooves or including a more proximal compartment, are possible modifications of the current design.
ii. Our results are based on unmyelinated axons. Co-culture with glia cells that may play a role in AxD is possible in the presented microfluidic device and the time course and morphological changes may be different under co-culture conditions. These studies are of high relevance to the field, but go beyond the scope of the present study.
iii. The observed effects of AxD in hemorrhagic stroke within this study were based on hemin toxicity, and we cannot exclude that other hemolysis products such as thrombin or bilirubin have different effects. Additional studies should investigate differences of hemolysis products to increase our understanding of the mechanisms of AxD in hemorrhagic stroke.
iv. The overall CNN performance may be further improved with more general inputs. For example, the segmentation of fragment pixels cannot be conducted accurately based on a single image at a specific time point as the whole process of AxD, ultimately resulting in the disintegration of the axons (i.e., the generation of axonal fragments), needs to be considered. CNNs using 3D convolutions could, in principle, perform a segmentation over an entire time-lapse recording and model temporal dependencies. However, we decided against the 3D approach, as it severely restricts general applicability due to its greatly increased effort to label suitable time series for training. In this context, the identification of the images that will yield the best results is crucial to effectively reduce labeling costs, which we have previously described using an active learning method (Grüning, P. et al., 2020).

## Conclusion

In combination with an advanced microfluidic device, the EntireAxon deep learning tool expands our possibilities to track AxD by detecting axons, axonal swellings, and axonal fragments. We further identified four morphological patterns of AxD, i.e., granular, retraction, swelling, and transport degeneration, under pathophysiological conditions in the context of hemorrhagic stroke. This approach will help to tackle the complex processes of AxD and may significantly enhance our understanding of AxD in health and disease to develop novel therapeutic strategies for brain diseases.

## Methods

Chemicals and reagents are listed in Tables S4-5.

### Study design

#### Sample size

Six mice. We did not perform a priori power analysis as this was an exploratory study. We did not change the number of the mice during the course of the study.

#### Data inclusion/exclusion criteria

Recordings that did not have any technical flaws, such as shifting of the microfluidic device in x, y, or z-axis were included. Recordings with minor x and y-axis shifts that we were able to correct by post-recording alignment (see **Image preprocessing**) were included. All data were processed using the same settings. The training and validation images for the deep learning tool were chosen to represent the testing data as best as possible.

#### Outliers

No outliers have been excluded in the study.

#### Selection of endpoints

Endpoints were the area of the axons, axonal swellings, and axonal fragments, respectively.

#### Replicates

Each individual mouse counted as a biological replicate (N = 6 biological replicate per experiment). Four different microfluidic units have been used for four experimental conditions (0, 50, 100, 200 μM hemin) per biological replicate.

#### Research objectives

The research objective was to examine the progression of axonal degeneration in primary cortical neurons upon hemin exposure. Therefore, a microfluidic device and deep learning tool to increase the experimental yield and to enable unbiased automatic analysis was developed. Our pre-specified hypothesis was to detect a concentration-dependent effect of hemin on axons. Our suggested hypothesis after conducting time-lapse recording was that there are four morphological patterns of axonal degeneration and that those depend on the severity of axonal degeneration by different hemin concentrations.

#### Research subjects or units of investigation

We employed primary cortical neurons from Crl:CD1 (ICR) Swiss outbred mice.

#### Experimental design

Randomized controlled laboratory experiment with four different concentrations of hemin treatment to induce and record axonal degeneration by time-lapse microscopy and further validation by fluorescence microscopy.

#### Randomization

Microfluidic units have randomly been assigned to one of the four experimental conditions (0, 50, 100, 200 μM hemin).

#### Blinding

The experimenter was not blinded when axons were treated with different hemin concentrations. The actual analysis was objective as being conducted solely by the deep learning tool.

### Fabrication of an enhanced throughput microfluidic device based on soft lithographic replica molding

Thirty-two wells were milled in a polymethyl methacrylate (PMMA) plate of the size of a conventional cell culture plate (**Fig. 1A and Supplementary Fig. S1**) using a universal milling machine (Mikron WF21C, Mikron Holding AG) with a 1 mm triple tooth cutter (HSS-CO8 Type N, Holex) at a precision of 0.01 mm. During the milling procedure, we applied a half-synthetic cooling lubricant (Opta Cool 600 HS, Wisura GmbH) on a mineral base to reduce the debris. Additionally, we milled screw holes in the intermediate spaces between each microfluidic unit to later detach the PMMA from the negative casting mold. To remove debris, we washed the PMMA plate by sonication (Sonicator Elmasonic S, Elma Schmidbauer GmbH) at room temperature for 30 minutes. Next, we lasered the microgrooves on the PMMA plate to connect both milled compartments of each individual microfluidic unit by using an Excimerlaser (Excistar XS 193 nm, Coherent). The PMMA plate was then washed again by sonication at room temperature for 30 minutes.

Polydimethylsiloxane (PDMS) was prepared in a 1:10 ratio and mixed properly before inducing vacuum at 0.5 Torr in a vacuum desiccator (Jeio Tech VDC-31) for 30 minutes. After the PDMS was poured into an empty aluminum basin to cover the ground, we applied vacuum at 0.5 Torr for 30 minutes to remove air bubbles. The PDMS was cured at room temperature for 48 hours. We put the PMMA plate on top of the PDMS ground with the milled and lasered structures showing upwards. Half of each well of the microfluidic units was filled with PDMS before curing at room temperature for 48 hours. We mixed the epoxy solution in a 1:1 ratio and poured it over the microfluidic device to cover its surface by at least 1 cm. Vacuum was applied at 0.5 Torr for 10 minutes to remove all air bubbles located above the channel side of the microfluidic device. The epoxy was cured at room temperature for a minimum of 2 hours. We subsequently detached the epoxy from the PMMA plate via a metallic block that consisted of screw holes in the intermediate spaces between the individual systems. The epoxy represented a negative casting mold to produce the microfluidic devices using PDMS.

PDMS was prepared as described above. We poured the PDMS into the negative epoxy casting mold and applied vacuum at 0.5 Torr for 30 minutes. The liquid PDMS was cured at 75 °C for 2 hours to induce the polymerization. We peeled the microfluidic devices from the casting mold and punched the wells with an 8 mm biopsy punch (DocCheck Shop GmbH) to ensure a sufficient amount of medium for cell culture. We cleaned customized 115 x 78 x 1 mm glass slides by sonication (Sonicator Elmasonic S, Elma Schmidbauer GmbH) and subsequently cleaned them by ethanol before plasma treatment (High Power Expanded Plasma Cleaner, Harrick Plasma). Plasma was applied at 45 W and 0.5 Torr for 2 minutes to activate the silanol groups of the glass slides and the microfluidic devices enabling firm attachment.

We washed the microfluidic devices with ethanol and then twice with distilled water to remove any debris. After aspirating the distilled water, except from the inside of the compartments, 0.1 mg/mL of poly-d-lysine solution in 0.02 M borate buffer (0.25 % (w/v) borate acid, 0.38 % (w/v) sodium tetraborate in distilled water, pH 8.5) was used for coating at 4 °C overnight. We aspirated the poly-d-lysine the next morning, not removing it from the compartments, and added 50 μg/mL of laminin as a second coating surface for incubation at 4 °C overnight. At the day of neuron isolation, the microfluidic devices were washed twice with pre-warmed medium after aspirating the laminin. Immediately prior to cell seeding, we aspirated the medium from the wells without removing it from the compartments.

### Experimental animals

Crl:CD1 (ICR) Swiss outbred mice (Charles River) were used. The animals were kept at 20-22 °C, 30-70 % humidity in a 12-hour/12-hour light/dark cycle and were fed a standard chow diet (Altromin Spezialfutter GmbH) *ad libitum*. Animal experiments followed the protocol of the “NIH Guide for the care and use of laboratory animals” and were approved by the Schleswig-Holstein Ministry for Energy Transition, Agriculture, Environment, Nature and Digitalization (under the prospective contingent animal license number 2017-07-06 Zille).

### Isolation and culture of primary cortical neurons

We isolated primary cortical neurons from murine E14 embryos after decapitation as previously described (Zille et al., 2017). We seeded the neurons at a density of 10,000 cells/mm^2^ in 5 μL MEM+Glutamax medium into one compartment (soma compartment) of each microfluidic unit of the device. The cells were allowed to adhere at 37 °C for 30 minutes. In order to promote directional axon growth into the other compartment (axonal compartment) by medium microflux, 150 μL of MEM+Glutamax medium were applied to the well of the soma compartment, while 100 μL were added to the well of the axonal compartment (**Fig. 1B**). Neurons were cultured at 37 °C in a humidified 5 % CO2 atmosphere. The next day, we changed from MEM+Glutamax medium to Neurobasal Plus Medium containing 2 % B-27 Plus Supplement, 1 mM sodium pyruvate and 1 % penicillin/streptomycin. The volume differences among the wells ensured the microflux for the directional axonal growth over the following days.

### Immunofluorescence

Soma and axonal compartments in the microfluidic units were fixed at room temperature for 1 hour in 4 % formaldehyde solution in phosphate buffered saline (PBS). They were washed twice with PBS and permeabilized with blocking solution (2 % BSA, 0.5% Triton-X-100 and 1x PBS) at room temperature for 1 hour. We incubated the neurons/axons on both compartments with primary antibodies against synaptophysin (1:250) and MAP2 (1:4000) at 4 °C overnight. The next day, both compartments were washed three times with PBS and incubated with the secondary antibodies anti-mouse Alexa Fluor 546 (1:500) and anti-rabbit Alexa Fluor 488 (1:500) at room temperature for 1 hour. After washing three times with PBS, both compartments were incubated with DAPI (1 μg/mL) for nuclear counterstaining at room temperature for 10 minutes. Both compartments were washed three times with PBS prior to fluorescence microscopy. An Olympus IX81 time-lapse microscope (Olympus Deutschland GmbH) with a 10X objective (0.3 NA Ph1) and camera F-View soft Imaging system was used at room temperature. Images were acquired with Cell^M^ software (Olympus Deutschland GmbH) and further processed via ImageJ (see **Image preprocessing**).

### Selection of microfluidic units for hemin treatment and time-lapse recording

At six or seven days in culture, microfluidic devices were considered for recording if they met the following inclusion criteria: i) axon growth through at least 80 % of all microgrooves and ii) axon length of at least 150 μm from the end of the microgrooves. All included microfluidic units were randomly assigned to the experimental conditions.

### Time-lapse recording of axonal degeneration

Axons were treated with 0 (vehicle), 50, 100, and 200 μM hemin. For the treatment, the medium was removed from the wells of the microfluidic units; hemin was diluted in the collected media and added back to the respective wells. The media volume between the two wells was equalized during the treatment to prevent any microflux. All microfluidic units were recorded immediately after each other. We started the recordings at 1 hour after treatment to allow for the adjustment of the well plates to the humidity of the incubation chamber of the microscope and the setup of the recording positions. We recorded AxD in Neurobasal Plus Medium containing 2 % B-27 Plus Supplement, 1 mM sodium pyruvate and 1 % penicillin/streptomycin with a 30-minutes interval for 24 hours using an Olympus IX81 time-lapse microscope (see **Immunofluorescence**) at 37 °C, 5 % CO2 and 65 % humidity.

### Live cell fluorescent staining

To evaluate axonal vitality, we washed the axonal compartment once with PBS and incubated the axonal compartment with calcein AM (4 μM) in PBS for 30 minutes at 37 °C at the end of the time-lapse recording or in 4-hour intervals upon hemin treatment. An Olympus IX81 time-lapse microscope (see **Immunofluorescence**) was used to record the respective images at 37 °C, 5 % CO_2_ and 65 % humidity.

### Training of the EntireAxon CNN for the segmentation of phase-contrast microscopic images

We trained the EntireAxon CNN for the image-wise semantic segmentation of AxD features in a supervised manner (**Fig. 2B**). To this end, we adapted a standard u-net with ResNet-50 encoder (Ronneberger et al., 2015) to automatically determine the class probability for each pixel of an input image. Our segmentation aimed to classify each pixel of a microscopic image of a time-lapse recording into one of four classes: ‘background’, ‘axon’, ‘axonal swelling’, and ‘axonal fragment’. For the training dataset, we selected 33 images and created corresponding image labels (masks) using GIMP (v.2.10.14, RRID:SCR_003182). For each image, a label image with the same height and width was created, in which each pixel value denotes a pixel class. Specifically, the classes ‘background’, ‘axon’, ‘axonal swelling’, and ‘axonal fragment’ had the values 0, 1, 2, and 3, respectively. For each pixel of the input image, we retained 4 values that reflect the probability distribution of the pixel over the four classes. We assigned each pixel the most probable class to create a segmentation map. During training, the CNN observed an input image, produced an output and compared this output to the label. The weights of the network were adapted via backpropagation so that the output better fitted the label. The weight changes were derived from a pixelwise loss function, i.e. the cross-entropy loss:

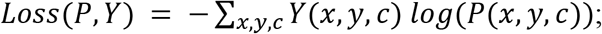

with *P*(*x, y, c*) *and Y*(*x, y, c*) being the probability of class c at pixel (*x, y*) for the prediction and ground truth of the network, respectively.

We trained a mean ensemble consisting of eight neural networks for 180 epochs using the Adam optimizer, a batch size of four and a learning rate of 0.001 that decreased by a factor of ten after every 60 epochs. The input images were standardized by the image-net mean and standard deviation (Deng et al., 2009). For data augmentation, we used random cropping (size 512 x 512), image flipping along the horizontal axis and rotation by a random angle between −90° and +90°.

### Validation of the EntireAxon CNN compared to human experts

To measure how well the EntireAxon CNN segments unknown images (**Fig. 2C**), we used a second validation set comprising eight images that were labeled by three human experts (A. Palumbo, S.K.L., L.E.H.). Importantly, the EntireAxon CNN did not update its parameters during training to fit the validation set, but only used the training set.

For each image, the EntireAxon CNN inferred a segmentation. We generated a binary mask from the prediction of the network, where 1 denotes the respective class and 0 all other classes. We computed a binary label mask in the same manner. We counted the true positive (TP), false positive (FP), and false negative (FN) pixels and computed the recall (sensitivity) and precision (Forman and Scholz, 2010):

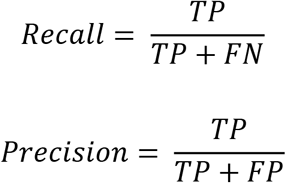

Recall and precision were calculated for each class separately on each validation image. The mean recall and precision over all eight validation images were determined subsequently.

A mean of 96.42 % of pixels in the axonal images were ‘background’ pixels, while only 2.77 % represented the class ‘axon’, 0.58 % ‘axonal swelling’, and 0.23 % ‘axonal fragment’ pixels. This reflects a challenging degree of class imbalance, where the probability of having any positives for a class in a validation image is low. Thus, we did not use the computed recall and precision of the individual images or the mean recall and precision to compute the mean F1 score, i.e., the harmonic mean of recall and precision. This has been shown to lead to bias, especially when a high degree of class imbalance is present in the dataset (Forman and Scholz, 2010) as it may result in undefined values for an image for recall (due to the absence of TP), precision (in case the CNN does not recognize the few positives), and F1 score (in case either recall or precision are undefined). To avoid bias, we computed the total TP, FP, and FN of all validation images from which we calculated the mean F1 score (Forman and Scholz, 2010):

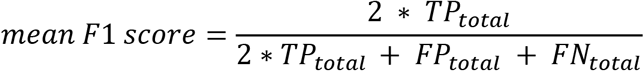

In addition, we computed a consensus label between human expert 1 and 2, 1 and 3 as well as 2 and 3 and compared the EntireAxon CNN versus the remaining expert (human expert 3, 2, and 1, respectively) to the consensus labels. Mean F1 scores for all classes were computed as described above.

### Image preprocessing

Prior to the analysis of AxD after hemin exposure, we preprocessed the time-lapse recordings in ImageJ (v1.52a, RRID: RRID:SCR_003070) using a custom-written macro. Specifically, each individual recording was converted from a 16-bit into an 8-bit recording to make it compatible with the ImageNet (8-bit) pre-trained ResNet-50. The recording was aligned automatically with the ImageJ plug-in “Linear Stack Alignment with SIFT” as described previously (Lowe, 2004). The following settings were used: initial Gaussian blur of 1.6 pixel, 3 steps per scale octave, minimum image size of 64 pixel, maximum image size of 1024 pixel, feature descriptor size of 4, 8 feature descriptor orientation bins, closest/next closest ratio of 0.92, maximal alignment error of 25 pixel; inlier ratio of 0.05, expected transformation as rigid, “interpolate” and “show info” checked. Black edges appearing on the recording after alignment were cropped.

### AxD analysis using the EntireAxon CNN

All recordings of AxD after hemin exposure were automatically analyzed by the trained EntireAxon CNN, which classified each pixel as one of the four different classes ‘background’, ‘axon’, ‘axonal swelling’, and ‘axonal fragment. For each experimental condition (i.e. hemin concentration), the sum percentage of all pixels per class on all images of that experimental day were added at each time point (‘Axont_1.5–24h_, Axonal swelling_t1.5-24h_, Axonal fragment_t1.5-24h_). To determine the changes for the classes ‘axon’, ‘axonal swelling’, and ‘axonal fragment’ over time, we calculated the sum percentage of pixels for all given time points (t_i_ with I = 1.5 to 24 hours) of the corresponding class over the sum of the pixels of all three classes at baseline:

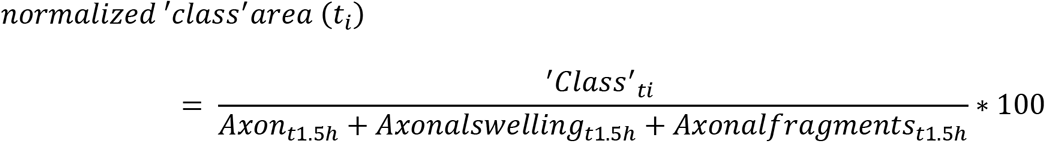

### Classification of the morphological patterns of AxD using attention-based RNN

We used the segmentation videos derived from the original microscopic images using the CNN to identify four morphological patterns of AxD: granular, retraction, swelling, and transport degeneration (**Fig 7A**). To reduce the dimensions of the input, the segmentation video was converted into a series of normalized histograms (*H*), one for each (time) frame. Thus, the RNN did not operate on the microscopic images directly, but rather on more efficient representations of the data. To compute a histogram for a frame ti, we compared the pixels of the frames t_i_ and t_i+1_. Each pixel was assigned into one of 16 classes that consisted of pairs (*c*_1_, *c*_2_) ∈ {0,1,2,3}^2^ of the four segmentation classes (i.e., four times four possible configurations, 16 class pairs). For example, the class (background, axon) means that in frame t_i_, the pixel was classified as background, while in frame t_i+1_, it was an axon pixel. For T time steps, we therefore computed T-1 histograms. *H*_0_(*t_i_*, (*c*_1_, *c*_2_)) is the number of pixels that belong to class *c*_1_ at time-frame and that belong to *c*_2_ at time-frame *t*_*i*+1_. Additionally, we normalized each histogram to sum up to 1 (i.e. we divided by the sum over all pairs):

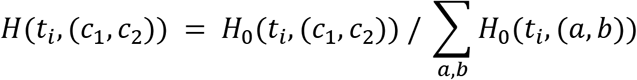

Of note, the histograms were computed over small patches (height and width < 90 pixels) during training and during inference on windows of size 32×32 pixels.

We used an encoder-decoder RNN with attention (Bahdanau et al., 2016). The encoder *f_enc_* consisted of a gated recurrent unit (GRU) that obtained the histogram time sequence *H* as input. The encoder computed the hidden representation of the histograms:

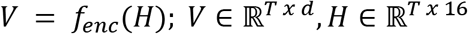

For our purpose, we used an architecture that was able to base the decision for a degeneration class on the previous class predictions. To this end, the output 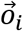 was computed iteratively in C+1 steps as a sum of the previous output and the output of the decoder *f_dec_*:

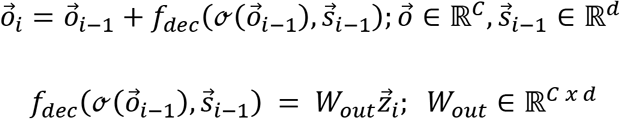

C is the number of degeneration classes (4) and d is the hidden dimension (we used 256); *i* = 1, *1*, C + 1. 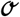 is the sigmoid function. The decoder employed a GRU that depended on the context vector 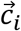 and the hidden state vector 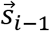:

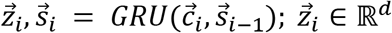

The entries of the initial hidden vector 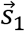 were all zero. The context vector is a weighted sum of the encoder representations. At each iteration, these weights can change, enabling the network to focus on different time-steps. We assumed that a specific pattern of degeneration happened only in a limited number of time frames that were fewer than the whole input video. The weights depended on the current state of the decoder and the current output:

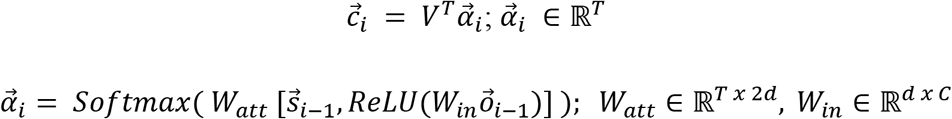

Here, 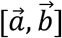 is the concatenation of two vectors. The final output *y* is normalized by the sigmoid function:

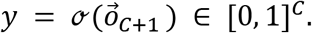

Apart from the weights used by the GRUs, *W_in_, W_att_*, and *W_out_* are learnable weights.

The EntireAxon RNN was trained with 162 images for 60 epochs using the lamb optimizer (You et al., 2020) with a batch size of 128. We used a learning rate of 0.01 that was reduced by a factor of ten every 15 epochs and an additional weight decay of 0.0001. The two GRUs (encoder and decoder) contained three layers, and we used dropout with a p-value of 0.9. To increase the RNN robustness against varying axon thickness, we also added eroded versions of the segmentation data using a cross-shape as kernel with the sizes three, five, and seven. Accordingly, each image existed six times in the dataset: three eroded versions and three unchanged copies, to keep a 50 % chance of having the original image for training.

### RNN cluster analysis

The unnormalized class output 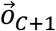 was computed by the matrix-vector product 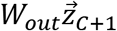. Where 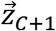 was a 256-dimensional vector representation of the input sample, computed by the model. For the classes to be linearly separable, the vector representations of each class needed to be close to each other in the 256-dimensional space. To visualize the relationships of the specific samples, we employed t-distributed stochastic neighborhood embedding (T-SNE) to compute a 2-dimensional representation of the high-dimensional data.

### Ten-fold cross-validation of the RNN

To validate the RNN, we used ten-fold cross-validation (Hastie et al., 2009). The dataset *S* was divided into 10 subsets, ensuring that each subset included at least one sample of each class: 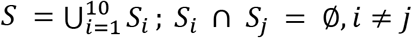. We trained ten models for *i*=1,…,10 on *Train_i_* = *S* / *S_i_* and test them on *Test_i_*=*S_i_*. Subsequently, we combined and evaluated all test samples 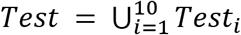. Mean recall, precision, and F1 score were determined as described above.

### Analysis of morphological pattern of AxD using the EntireAxon RNN

All AxD segmentations after hemin exposure were automatically analyzed with the trained EntireAxon RNN, which predicted the occurrence of the four morphological patterns of AxD in a pixel-wise manner. Of note, a pixel can be predicted to belong to 0, 1 or multiple morphological patterns. Only pixels previously identified as degenerated over time were considered by applying a ‘fragmentation mask’ that included all no-background pixels that changed to either background or fragment during the recording time.

For each experimental condition (i.e., hemin concentration), the percentage of the occurrence of each morphological pattern was calculated as the sum of all pixels per morphological pattern on all images of that experimental day divided by the ‘fragmentation mask’ as follows:

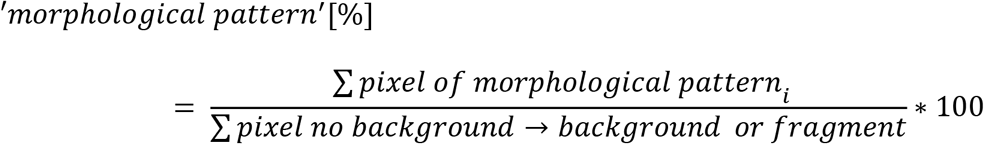

### Statistical analysis

Six biological replicates for each concentration were employed in each experiment to assess hemin-induced AxD. We did not perform a priori power analysis as this was an exploratory study. Normality was evaluated with the Kolmogorov-Smirnov test, variance homogeneity using the Levené test, and sphericity by the Mauchly test. When the data were normally distributed and variance homogeneity was met, one-way ANOVA followed by the Bonferroni post hoc test was performed. In case the data were not normally distributed, the Kruskal-Wallis test was performed for multiple comparisons of independent groups followed by the post hoc Mann-Whitney U test with α-correction according to Bonferroni to adjust for the inflation of type I error due to multiple testing. For the repeated testing with covariates, a repeated measures ANOVA was performed with Greenhouse-Geisser adjustment if sphericity was not given. Linear regressions were performed for AxD patterns. Data are represented as mean ± 95 % confidence interval (CI) except for the nonparametric data of the AUC for axonal fragments, where medians are given. A value of *P* < 0.05 was considered statistically significant. For the Kruskal-Wallis test followed by Mann-Whitney U, *P* =0.05/k was used, with k as the number of single hypotheses. K = 3 for AUC analyses (comparison of three different concentrations of hemin vs. 0 μM hemin), thus α = 0.0167 was considered statistically significant. K = 6 for the comparison of the linear regression slopes (comparison of the four AxD patterns against each other), thus α = 0.0083 was considered statistically significant. The detailed statistical analyses can be found in **Tables S1-3**. All statistical analyses were performed with IBM SPSS version 23 (RRID:SCR_002865), except linear regressions that were performed with GraphPad Prism version 8 (RRID:SCR_002798).

## List of abbreviations

AxD: axonal degeneration
CNN: convolutional neural network
FP: false positive
FN: false negative
GRU: gated recurrent unit
PBS: phosphate buffered saline
PDMS: polydimethylsiloxane
PMMA: polymethyl methacrylate
RNN: recurrent neural network
TP: true positive

## Declarations

## Acknowledgments

This research was supported by a Fraunhofer MEF grant of the Fraunhofer Society (project number 600199) and by the Joachim Herz Stiftung (project number 850022) to M.Z. We would like to thank Sebastian Kärst for his technical assistance regarding the choice of materials for the prototype of the microfluidic device. We would like to thank Dennis Wendt for his help in designing the blueprint of the prototype of the microfluidic device.

## Authors’ contributions

M.Z. designed the experiments. A. Palumbo, A. Pabst and M.Z. designed the device. A. Palumbo, A. Pabst, S.P., R.S., C.K., and N.K. carried out the fabrication of the device. S.K.L. performed the immunostaining of the somata and axons and analyzed the respective data. P.G. and A.M.M. developed the deep learning tool, P.G., L.E.H., and L.B. developed the algorithms to retrieve the output. A. Palumbo, S.K.L. and L.E.H. labeled the images for the deep learning training and validation. A. Palumbo and C.F. performed the time-lapse recordings of AxD. A. Palumbo conducted the live cell imaging, the determination of the morphological patterns of AxD and analyzed the data for the respective experiments. A. Palumbo, P.G., A.M.M., J.B., and M.Z. discussed and interpreted the data. M.Z. performed the statistical analysis. A. Palumbo, M.I. and M.Z. performed the graphical artwork. A. Palumbo, P.G., and M.Z wrote the manuscript. All authors discussed and commented on the final version of the manuscript.

## Ethics approval

Animal experiments followed the protocol of the “NIH Guide for the care and use of laboratory animals” and were approved by the Schleswig-Holstein Ministry for Energy Transition, Agriculture, Environment, Nature and Digitalization (under the prospective contingent animal license number 2017-07-06 Zille).

## Data availability statement

All data needed to evaluate the conclusions in the paper are present in the paper and/or the **Supporting information**. The time-lapse data and code are available upon reasonable request to the corresponding authors. We plan to launch a website to enable other researchers to use the tool.

## References

Bahdanau, D., K. Cho, and Y. Bengio. 2016. Neural Machine Translation by Jointly Learning to Align and Translate. arXiv:1409.0473 [cs, stat].

Becker, T., and A. Madany. 2012. Morphology-based Features for Adaptive Mitosis Detection of In Vitro Stem Cell Tracking Data. Methods Inf Med. 51:449–456. doi:10.3414/ME11-02-0038.

Beirowski, B., R. Adalbert, D. Wagner, D.S. Grumme, K. Addicks, R.R. Ribchester, and M.P. Coleman. 2005. The progressive nature of Wallerian degeneration in wild-type and slow Wallerian degeneration (WldS) nerves. BMC Neurosci. 6:6. doi:10.1186/1471-2202-6-6.

Biffi, E. 2015. Microfluidic and Compartmentalized Platforms for Neurobiological Research.

Bishop, D.L., T. Misgeld, M.K. Walsh, W.-B. Gan, and J.W. Lichtman. 2004. Axon Branch Removal at Developing Synapses by Axosome Shedding. Neuron. 44:651–661. doi:10.1016/j.neuron.2004.10.026.

Cavanagh, J.B. 1979. The “dying back” process. A common denominator in many naturally occurring and toxic neuropathies. Arch. Pathol. Lab. Med. 103:659–664.

Chen, J., and R.F. Regan. 2004. Heme oxygenase-2 gene deletion increases astrocyte vulnerability to hemin. Biochemical and Biophysical Research Communications. 318:88–94. doi:10.1016/j.bbrc.2004.03.187.

Chen, X., X. Chen, Y. Chen, M. Xu, T. Yu, and J. Li. 2018. The Impact of Intracerebral Hemorrhage on the Progression of White Matter Hyperintensity. Front. Hum. Neurosci. 12:471. doi:10.3389/fnhum.2018.00471.

Coleman, M.P. 2005. Axon degeneration mechanisms: commonality amid diversity. Nat Rev Neurosci. 6:889–898. doi:10.1038/nrn1788.

Cui, Y., X. Jin, D.-J. Choi, J.Y. Choi, H.S. Kim, D.H. Hwang, and B.G. Kim. 2020. Axonal degeneration in an in vitro model of ischemic white matter injury. Neurobiology of Disease. 134:104672. doi:10.1016/j.nbd.2019.104672.

Deng, J., W. Dong, R. Socher, L.-J. Li, Kai Li, and Li Fei-Fei. 2009. ImageNet: A large-scale hierarchical image database. In 2009 IEEE Conference on Computer Vision and Pattern Recognition. IEEE, Miami, FL. 248–255.

Dietterich, T.G. 2000. Ensemble Methods in Machine Learning. In Multiple Classifier Systems. Springer Berlin Heidelberg, Berlin, Heidelberg. 1–15.

Forman, G., and M. Scholz. 2010. Apples-to-apples in cross-validation studies: pitfalls in classifier performance measurement. SIGKDD Explor. Newsl. 12:49. doi:10.1145/1882471.1882479.

Grüning, P., Palumbo, A., Zille, M., Barth, E., and Madany Mamlouk, Amir. 2020. A task-dependent active learning method for axon segmentation with CNNs. Proc AUTOMED. (1). doi:10.18416/AUTOMED.2020.

Hastie, T., R. Tibshirani, and J. Friedman. 2009. The Elements of Statistical Learning. Springer New York, New York, NY.

He, K., X. Zhang, S. Ren, and J. Sun. 2015. Deep Residual Learning for Image Recognition. arXiv:1512.03385 [cs].

Hemorrhagic Stroke Academia Industry (HEADS) Roundtable Participants. 2018. Basic and Translational Research in Intracerebral Hemorrhage: Limitations, Priorities, and Recommendations. Stroke. 49:1308–1314. doi:10.1161/STROKEAHA.117.019539.

Ho, S.-Y., C.-Y. Chao, H.-L. Huang, T.-W. Chiu, P. Charoenkwan, and E. Hwang. 2011. NeurphologyJ: An automatic neuronal morphology quantification method and its application in pharmacological discovery. BMC Bioinformatics. 12:230. doi:10.1186/1471-2105-12-230.

Hosmane, S., I.H. Yang, A. Ruffin, N. Thakor, and A. Venkatesan. 2010. Circular compartmentalized microfluidic platform: Study of axon–glia interactions. Lab Chip. 10:741. doi:10.1039/b918640a.

Huang, H.-K., C.-F. Chiu, C.-H. Kuo, Y.-C. Wu, N.N.Y. Chu, and P.-C. Chang. 2016. Mixture of deep CNN-based ensemble model for image retrieval. In 2016 IEEE 5th Global Conference on Consumer Electronics. IEEE, Kyoto, Japan. 1–2.

Kerschensteiner, M., M.E. Schwab, J.W. Lichtman, and T. Misgeld. 2005. In vivo imaging of axonal degeneration and regeneration in the injured spinal cord. Nat Med. 11:572–577. doi:10.1038/nm1229.

Li, Y., M. Yang, Z. Huang, X. Chen, M.T. Maloney, L. Zhu, J. Liu, Y. Yang, S. Du, X. Jiang, and J.Y. Wu. 2014. AxonQuant: A Microfluidic Chamber Culture-Coupled Algorithm That Allows High-Throughput Quantification of Axonal Damage. Neurosignals. 22:14–29. doi:10.1159/000358092.

Lingor, P., J.C. Koch, L. Tönges, and M. Bähr. 2012. Axonal degeneration as a therapeutic target in the CNS. Cell Tissue Res. 349:289–311. doi:10.1007/s00441-012-1362-3.

Lowe, D.G. 2004. Distinctive Image Features from Scale-Invariant Keypoints. International Journal of Computer Vision. 60:91–110. doi:10.1023/B:VISI.0000029664.99615.94.

Luo, L., and D.D.M. O’Leary. 2005. Axon Retraction and Degeneration in Development and Disease. Annu. Rev. Neurosci. 28:127–156. doi:10.1146/annurev.neuro.28.061604.135632.

Maor-Nof, M., E. Romi, H. Sar Shalom, V. Ulisse, C. Raanan, A. Nof, D. Leshkowitz, R. Lang, and A. Yaron. 2016. Axonal Degeneration Is Regulated by a Transcriptional Program that Coordinates Expression of Pro- and Anti-degenerative Factors. Neuron. 92:991–1006. doi:10.1016/j.neuron.2016.10.061.

Neukomm, L.J., and M.R. Freeman. 2014. Diverse cellular and molecular modes of axon degeneration. Trends in Cell Biology. 24:515–523. doi:10.1016/j.tcb.2014.04.003.

Nikić, I., D. Merkler, C. Sorbara, M. Brinkoetter, M. Kreutzfeldt, F.M. Bareyre, W. Brück, D. Bishop, T. Misgeld, and M. Kerschensteiner. 2011. A reversible form of axon damage in experimental autoimmune encephalomyelitis and multiple sclerosis. Nat Med. 17:495–499. doi:10.1038/nm.2324.

Nikolaev, A., T. McLaughlin, D.D.M. O’Leary, and M. Tessier-Lavigne. 2009. APP binds DR6 to trigger axon pruning and neuron death via distinct caspases. Nature. 457:981–989. doi:10.1038/nature07767.

Park, J.W., B. Vahidi, A.M. Taylor, S.W. Rhee, and N.L. Jeon. 2006. Microfluidic culture platform for neuroscience research. Nat Protoc. 1:2128–2136. doi:10.1038/nprot.2006.316.

Pease, S.E., and R.A. Segal. 2014. Preserve and protect: maintaining axons within functional circuits. Trends in Neurosciences. 37:572–582. doi:10.1016/j.tins.2014.07.007.

Pease-Raissi, S.E., M.F. Pazyra-Murphy, Y. Li, F. Wachter, Y. Fukuda, S.J. Fenstermacher, L.A. Barclay, G.H. Bird, L.D. Walensky, and R.A. Segal. 2017. Paclitaxel Reduces Axonal Bclw to Initiate IP3R1-Dependent Axon Degeneration. Neuron. 96:373–386.e6. doi:10.1016/j.neuron.2017.09.034.

Pool, M., J. Thiemann, A. Bar-Or, and A.E. Fournier. 2008. NeuriteTracer: A novel ImageJ plugin for automated quantification of neurite outgrowth. Journal of Neuroscience Methods. 168:134–139. doi:10.1016/j.jneumeth.2007.08.029.

Robinson, S.R., T.N. Dang, R. Dringen, and G.M. Bishop. 2009. Hemin toxicity: a preventable source of brain damage following hemorrhagic stroke. Redox Report. 14:228–235. doi:10.1179/135100009X12525712409931.

Ronneberger, O., P. Fischer, and T. Brox. 2015. U-Net: Convolutional Networks for Biomedical Image Segmentation. arXiv:1505.04597 [cs].

Salvadores, N., M. Sanhueza, P. Manque, and F.A. Court. 2017. Axonal Degeneration during Aging and Its Functional Role in Neurodegenerative Disorders. Front. Neurosci. 11:451. doi:10.3389/fnins.2017.00451.

Sasaki, Y., B.P.S. Vohra, F.E. Lund, and J. Milbrandt. 2009. Nicotinamide Mononucleotide Adenylyl Transferase-Mediated Axonal Protection Requires Enzymatic Activity But Not Increased Levels of Neuronal Nicotinamide Adenine Dinucleotide. Journal of Neuroscience. 29:5525–5535. doi:10.1523/JNEUROSCI.5469-08.2009.

Saxena, S., and P. Caroni. 2007. Mechanisms of axon degeneration: From development to disease. Progress in Neurobiology. 83:174–191. doi:10.1016/j.pneurobio.2007.07.007.

Tang-Schomer, M.D., V.E. Johnson, P.W. Baas, W. Stewart, and D.H. Smith. 2012. Partial interruption of axonal transport due to microtubule breakage accounts for the formation of periodic varicosities after traumatic axonal injury. Experimental Neurology. 233:364–372. doi:10.1016/j.expneurol.2011.10.030.

Tao, C., X. Hu, H. Li, and C. You. 2017. White Matter Injury after Intracerebral Hemorrhage: Pathophysiology and Therapeutic Strategies. Front. Hum. Neurosci. 11:422. doi:10.3389/fnhum.2017.00422.

Van Laar, V., B. Arnold, and S. Berman. 2019. Primary Embryonic Rat Cortical Neuronal Culture and Chronic Rotenone Treatment in Microfluidic Culture Devices. BIO-PROTOCOL. 9. doi:10.21769/BioProtoc.3192.

Venkatasubramanian, C., J.T. Kleinman, N.J. Fischbein, J. Olivot, A.D. Gean, I. Eyngorn, R.W. Snider, M. Mlynash, and C.A.C. Wijman. 2013. Natural History and Prognostic Value of Corticospinal Tract Wallerian Degeneration in Intracerebral Hemorrhage. JAHA. 2. doi:10.1161/JAHA.113.000090.

Vuola, A.O., S.U. Akram, and J. Kannala. 2019. Mask-RCNN and U-Net Ensembled for Nuclei Segmentation. In 2019 IEEE 16th International Symposium on Biomedical Imaging (ISBI 2019). IEEE, Venice, Italy. 208–212.

Wang, J.T., Z.A. Medress, and B.A. Barres. 2012. Axon degeneration: Molecular mechanisms of a self-destruction pathway. J Cell Biol. 196:7–18. doi:10.1083/jcb.201108111.

Yong, Y., K. Gamage, I. Cheng, K. Barford, A. Spano, B. Winckler, and C. Deppmann. 2019. p75NTR and DR6 Regulate Distinct Phases of Axon Degeneration Demarcated by Spheroid Rupture. J. Neurosci. 39:9503–9520. doi:10.1523/JNEUROSCI.1867-19.2019.

You, Y., J. Li, S. Reddi, J. Hseu, S. Kumar, S. Bhojanapalli, X. Song, J. Demmel, K. Keutzer, and C.-J. Hsieh. 2020. Large Batch Optimization for Deep Learning: Training BERT in 76 minutes. arXiv:1904.00962 [cs, stat].

Zille, M., S.S. Karuppagounder, Y. Chen, P.J. Gough, J. Bertin, J. Finger, T.A. Milner, E.A. Jonas, and R.R. Ratan. 2017. Neuronal Death After Hemorrhagic Stroke In Vitro and In Vivo Shares Features of Ferroptosis and Necroptosis. Stroke. 48:1033–1043. doi:10.1161/STROKEAHA.116.015609.

