## Supporting information for "Deep learning to decipher the progression and morphology of axonal degeneration"

**Running title:** EntireAxon deep learning of axonal degeneration

Alex Palumbo[1,2,3,*], Philipp Grüning[4], Svenja Kim Landt[1,2], Lara Eleen Heckmann[1,2], Luisa Bartram[1], Alessa Pabst[1,2], Charlotte Flory[1,2], Maulana Ikhsan[1,2,3], Sören Pietsch[1,2,5], Reinhard Schulz[6], Christopher Kren[7], Norbert Koop[7], Johannes Boltze[1,2,8], Amir Madany Mamlouk[4] and Marietta Zille[1,2,3,*]

[1] Fraunhofer Research and Development Center for Marine and Cellular Biotechnology, Fraunhofer Research Institution for Individualized and Cell-Based Medical Engineering, Lübeck, Germany

[2] Institute for Medical and Marine Biotechnology, University of Lübeck, Lübeck, Germany

[3] Institute for Experimental and Clinical Pharmacology and Toxicology, University of Lübeck, Lübeck, Germany

[4] Institute for Neuro- and Bioinformatics, University of Lübeck, Lübeck, Germany

[5] Department of Neonatology, Universitätsklinikum Leipzig, Leipzig, Germany

[6] Wissenschaftliche Werkstätten, University of Lübeck, Lübeck, Germany

[7] Medical Laser Center Lübeck GmbH, Lübeck, Germany

[8] School of Life Sciences, The University of Warwick, Gibbet Hill Campus, Coventry, United Kingdom

[*] Corresponding authors:

Alex Palumbo

Postal address: Fraunhofer Research and Development Center for Marine and Cellular Biotechnology, Fraunhofer Research Institution for Individualized and Cell-Based Medical Engineering, Mönkhofer Weg 239a, 23562, Lübeck, Germany

Marietta Zille

Postal address: Institute for Experimental and Clinical Pharmacology and Toxicology, University of Lübeck, Ratzeburger Allee 160, 23562, Lübeck, Germany

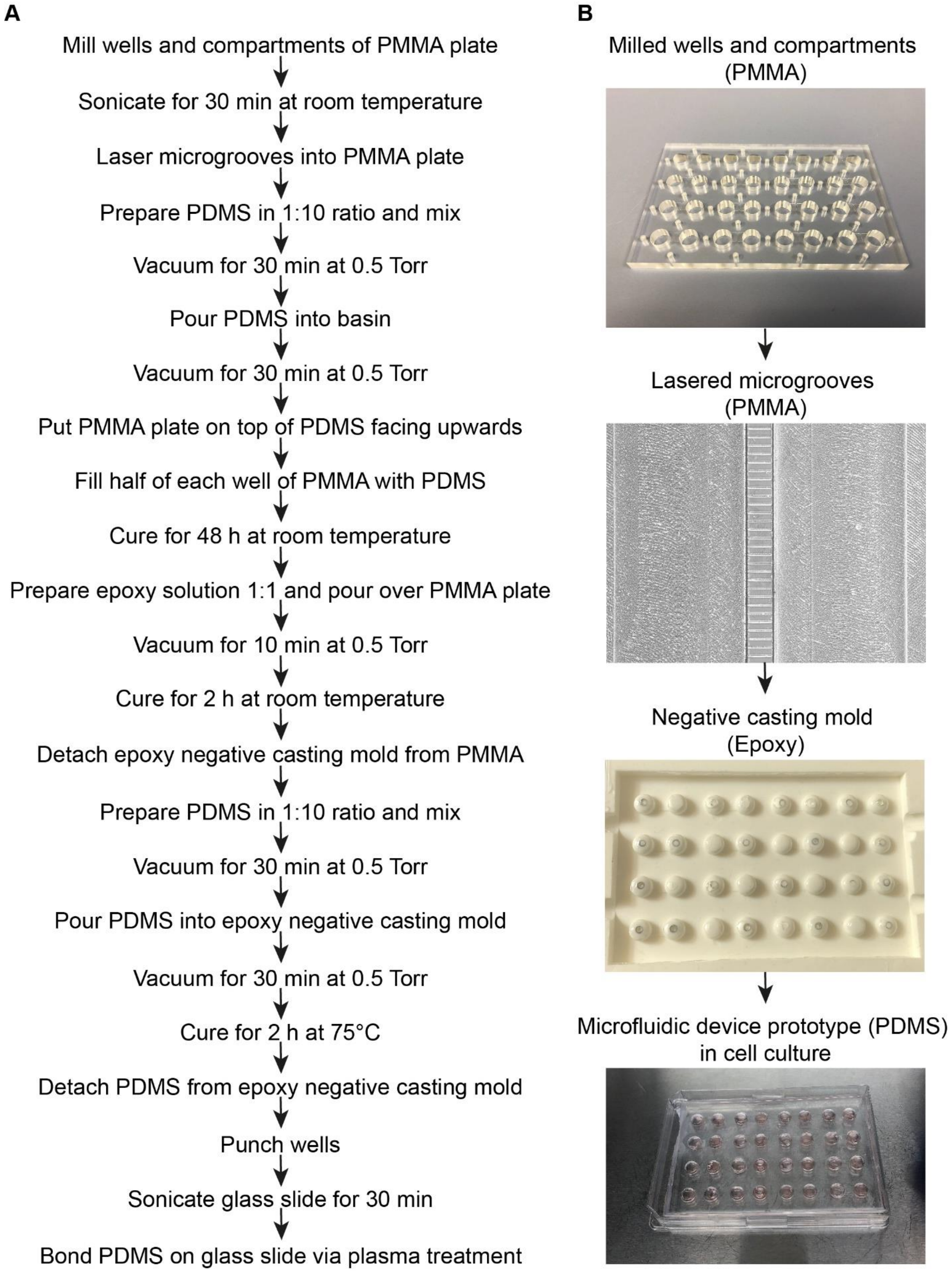


**Figure S1. Manufacturing of the microfluidic device for the enhanced throughput cultivation of axons. Related to Figure 1. (A)** Manufacturing process of the microfluidic device prototype. **(B)** Individual components of the production process. A milled and lasered polymethyl methacrylate (PMMA) plate was used as a positive imprint to develop a negative casting mold produced from epoxy. Polydimethylsiloxane (PDMS) was poured into the negative casting mold, bonded onto glass and applied in the cell culture.

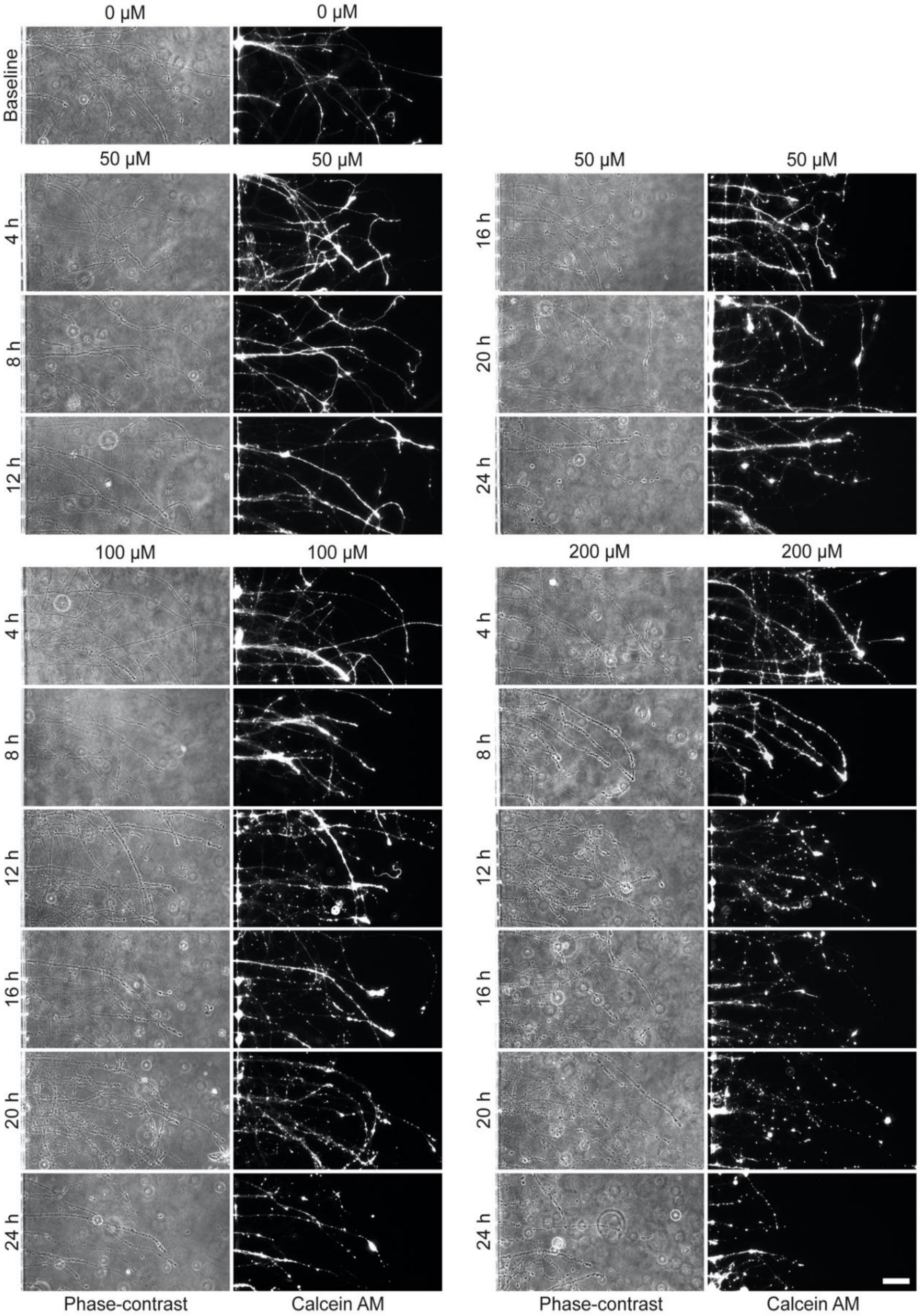


**Figure S2. Validation of the time course of axonal degeneration (AxD) by fluorescent live cell marker calein AM.** We determined the start and end of hemin-induced AxD by comparing the appearance of axonal swellings and axonal fragments in phase-contrast and fluorescence (calcein AM) microscopy. Compared to baseline (0 µM hemin), morphological hallmarks of AxD started to appear after 16 hours (50 µM), 12 hours (100 µM), and 8 hours (200 µM) and AxD was observed during the following 4 hours in all concentrations. N = 3 independent cultures of primary cortical neurons. Scale bar: 50 µm.

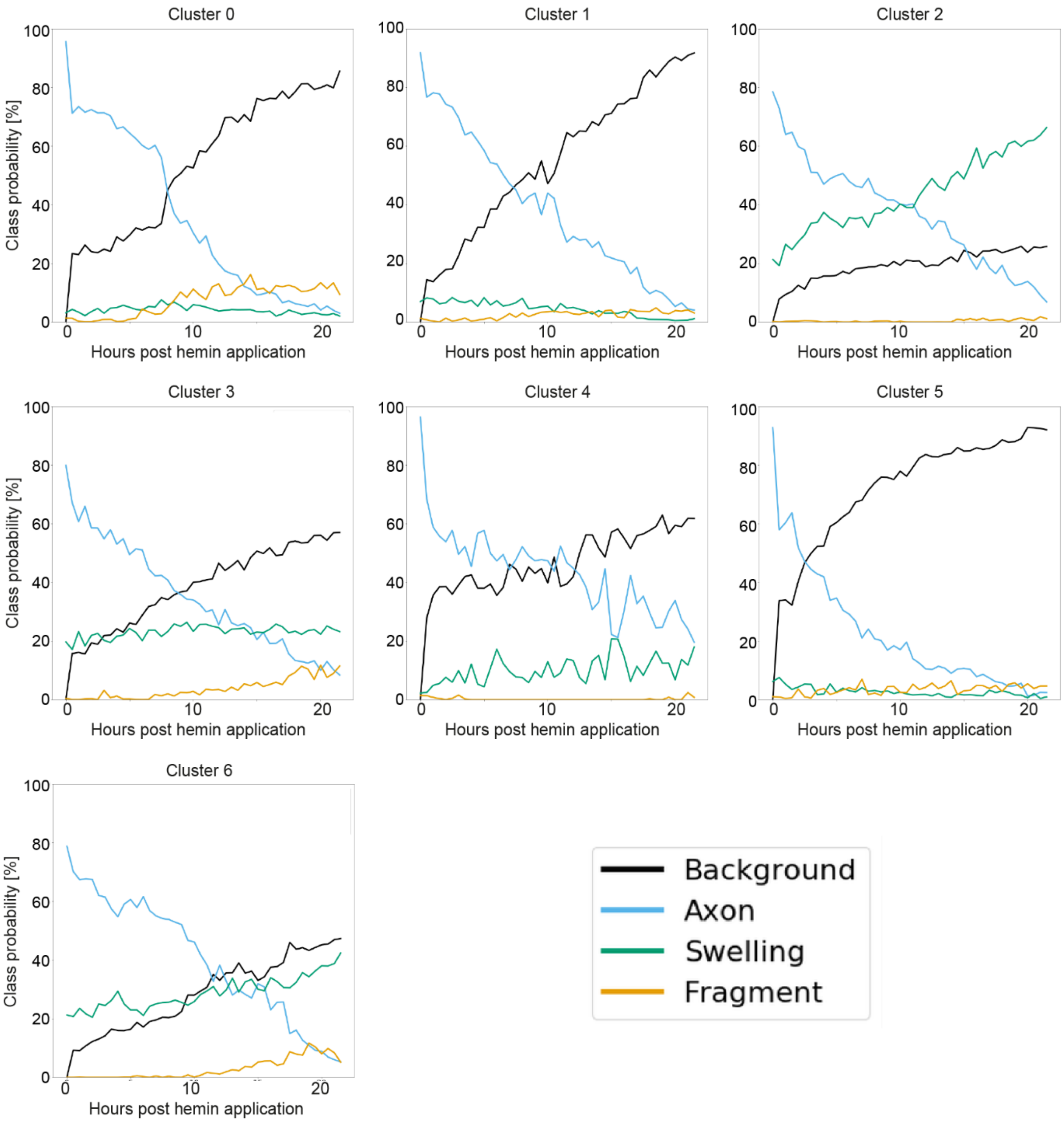


**Figure S3. Cluster analysis of the four morphological patterns of axonal degeneration (AxD) based on the changes in class segregation of the pixels.** For each time point in comparison to the previous time point, the RNN computed the probability of a change in class for each pixel of an image. Each cluster (0-6) was characterized by an idiosyncratic segmentation pattern over 24 hours.

**Table S1. Related to Fig. 3.** Time- and concentration-dependent AxD in an *in vitro* model of hemorrhagic stroke.

| Readout | Mauchly test | Omnibus Test | Posthoc Test |
| --- | --- | --- | --- |
| **Axon area** | [chi]2 is undefined because the number of repeated measurements is greater than the sample size; hence, sphericity was not assumed and the Greenhouse-Geisser correction was applied | one-way ANOVA with Greenhouse-Geisser correction ($\varepsilon$ = 0.045) across time*group: F(6.109) = 19.758, $P < 0.001$, partial-$\eta^2$ = 0.748 | Post hoc Bonferroni $P$ = 0.018 for 50 µM from 15 hours ($P < 0.001$ from 19 hours), $P$ = 0.040 for 100 µM from 14 hours ($P < 0.001$ from 18.5 hours), $P$ = 0.020 for 200 µM from 11.5 hours ($P < 0.001$ from 15 hours) vs. 0 µM |
| **Axonal swelling area** | | one-way ANOVA with Greenhouse-Geisser correction ($\varepsilon$ = 0.042) across time*group: F(5.703) = 3.201, $P$ = 0.013, partial-$\eta^2$ = 0.324 | Post hoc Bonferroni $P$ = 0.030 for 50 µM from 8 hours, $P$ = 0.019 for 100 µM from 6 hours, $P$ = 0.010 for 200 µM from 6 hours until 18.5 hours vs. 0 µM |
| **Axonal fragment area** | | one-way ANOVA with Greenhouse-Geisser correction ($\varepsilon$ = 0.026) across time*group: F(3.522) = 9.115, $P < 0.001$, partial-$\eta^2$ = 0.578 | Post hoc Bonferroni $P$ = 0.044 for 100 µM from 17 hours, $P$ = 0.037 for 200 µM from 9 hours vs. 0 µM |

**Table S2. Related to Fig. 4.** Area under the curve (AUC) analyses of hemin-mediated AxD.

| | **Kolmogorov-Smirnov test** | **Levené test** | **Omnibus Test** | **Posthoc Test** |
|---|---|---|---|---|
| **AUC Axon area** | Z = 0.093, *P* = 0.200 | F(3,20) = 0.116, *P* = 0.949 | one-way ANOVA F(3,20) = 8.547, *P* = 0.001, partial-$\eta^2$ = 0.562 | Post hoc Bonferroni *P* = 0.026 for 50 µM, *P* = 0.018 for 100 µM, and *P* < 0.001 for 200 µM vs. 0 µM |
| **AUC Axonal swelling area** | Z = 0.104, *P* = 0.200 | F(3,20) = 0.993, *P* = 0.416 | one-way ANOVA F(3,20) = 6.721, *P* = 0.003, partial-$\eta^2$ = 0.502 | Post hoc Bonferroni *P* = 0.012 for 50 µM, *P* = 0.005 for 100 µM, and *P* = 0.016 for 200 µM vs. 0 µM |
| **AUC Axonal fragment area** | Z = 0.245, *P* = 0.001 | F(3,20) = 5.800, *P* = 0.005 | Kruskal-Wallis Test [chi]2(3,N = 24) = 16.393, *P* = 0.001, $\eta^2$ = 0.713 | Post hoc Mann-Whitney test with Bonferroni correction at α = 0.0167, *P* = 0.037 for 50 µM, *P* = 0.004 for 100 µM and 200 µM vs. 0 µM |

**Table S3. Related to Fig. 8.** Comparison of the slopes of the linear regression of the four morphological patterns of AxD in an *in vitro* model of hemorrhagic stroke.

| | **Comparison of the slopes With Bonferroni correction at α = 0.0083 for multiple comparisons (six comparisons)** |
|---|---|
| **Granular degeneration vs. retraction degeneration** | F(1,44) = 6.971, *P* = 0.011, not significant at α = 0.008 |
| **Granular degeneration vs. swelling degeneration** | F(1,44) = 0.202, *P* = 0.655 |
| **Granular degeneration vs. transport degeneration** | F(1,44) = 8.865, *P* = 0.005, |
| **Retraction degeneration vs. swelling degeneration** | F(1,44) = 7.842, *P* = 0.0076, |
| **Retraction degeneration vs. transport degeneration** | F(1,44) = 0.406, *P* = 0.528, |
| **Swelling degeneration vs. transport degeneration** | F(1,44) = 9.074, *P* = 0.004, |

**Table S4. Chemicals and Reagents.**

| Product | Company | Catalogue number |
|---|---|---|
| B-27™ Plus Neuronal Culture System | Thermo Fisher Scientific | A3653401 |
| Boric acid | Merck | 925K12283365 |
| Bovin serum albumin | Sigma Aldrich | A9418 |
| Calcein AM | Santa Cruz | sc-203865 |
| Chicken trypsin inhibitor | Sigma Aldrich | T9253 |
| DAPI | Roche | 10236276001 |
| DNAse | Sigma Aldrich | D5025-15KU |
| Earle's balanced salt solution | Sigma Aldrich | E7510 |
| Epoxy solution Smooth-Cast 310/1 | KauPo | 09202-003-000002 |
| Ethanol 70% | Carl Roth | 9065.5 |
| Ethylenediaminetetraacetic acid | Carl Roth | 8043.3 |
| Fetal calf serum | Thermo Fisher Scientific | A15-151 |
| Hemin | Sigma Aldrich | H9039 |
| Horse serum | GE Health Life Sciences | SH30074.03 |
| Laminin | Sigma Aldrich | L2020 |
| L-cysteine | Carl Roth | 1693.2 |
| MEM, GlutaMAX™ Supplement | Thermo Fisher Scientific | 41090028 |
| Papain | Carl Roth | 8933.1 |
| Penicillin/Streptomycin | Biochrom | A2212 |
| Phosphate buffered saline | Thermo Fisher Scientific | 14200083 |
| Polydimethylsiloxane | Biesterfeld Spezialchemie GmbH | 5498840000 |
| Poly-d-lysine | Sigma Aldrich | P6407 |
| Polymethylmethacrylate | Kongsback | Customized |
| Sodium acetate | Carl Roth | X891.1 |
| Sodium bicarbonate | Sigma Aldrich | S5761 |
| Sodium pyruvate | Thermo Fisher Scientific | 11360070 |
| Sodium tetraborate | Merck | 1.06308.1000 |
| Triton-X-100 | Fluka | 93420 |

**Table S5. Antibodies.**

| Antibody | Company | Catalogue number | RRID |
|---|---|---|---|
| Polyclonal rabbit anti-microtubule-associated protein 2 (MAP2) | Abcam | ab32454 | AB_776174 |
| Monoclonal mouse anti-synaptophysin | Thermo Fisher Scientific | MA1-213 | AB_2723681 |
| Goat anti-mouse IgG (H+L) highly cross-adsorbed secondary antibody, Alexa Fluor 546 | Thermo Fisher Scientific | A-11030 | AB_2534089 |
| Goat anti-rabbit IgG (H+L) highly cross-adsorbed secondary antibody, Alexa Fluor 488 | Thermo Fisher Scientific | A-11034 | AB_2576217 |

### Supplementary Movies

**Video S1 (separate file). Time-lapse video of axons treated with 0 µM hemin.** Primary cortical axons were treated with vehicle (0 µM) for 24 hours and recorded by time-lapse microscopy in a 30-minutes interval and shown in 7 frames/second. Scale bar: 50 µm.
**Video S2 (separate file). Time-lapse video of axons treated with 50 µM hemin.** Primary cortical axons were treated with 50 µM hemin for 24 hours and recorded by time-lapse microscopy in a 30-minutes interval and shown in 7 frames/second. Scale bar: 50 µm.
**Video S3 (separate file). Time-lapse video of axons treated with 100 µM hemin.** Primary cortical axons were treated with 100 µM hemin for 24 hours and recorded by time-lapse microscopy in a 30-minutes interval and shown in 7 frames/second. Scale bar: 50 µm.
**Video S4 (separate file). Time-lapse video of axons treated with 200 µM hemin.** Primary cortical axons were treated with 200 µM hemin for 24 hours and recorded by time-lapse microscopy in a 30-minutes interval and shown in 7 frames/second. Scale bar: 50 µm.
**Video S5 (separate file). Time-lapse video of granular degeneration induced by hemin.** Granular degeneration of primary cortical axons treated with 200 µM hemin for 24 hours and recorded by time-lapse microscopy in a 30-minutes interval and shown in 7 frames/second. Scale bar: 20 µm.
**Video S6 (separate file). Time-lapse video of retraction degeneration induced by hemin.** Retraction degeneration of primary cortical axons treated with 200 µM hemin for 24 hours and recorded by time-lapse microscopy in a 30-minutes interval and shown in 7 frames/second. Scale bar: 20 µm.
**Video S7 (separate file). Time-lapse video of swelling degeneration induced by hemin.** Swelling degeneration of primary cortical axons treated with 200 µM hemin for 24 hours and recorded by time-lapse microscopy in a 30-minutes interval and shown in 7 frames/second. Scale bar: 20 µm.
**Video S8 (separate file). Time-lapse video of transport degeneration induced by hemin.** Transport degeneration of primary cortical axons treated with 200 µM hemin for 24 hours and recorded by time-lapse microscopy in a 30-minutes interval and shown in 7 frames/second. Scale bar: 20 µm.
**Video S9 (separate file). Time-lapse video of the segmentation by the recurrent neuronal networks of AxD induced by hemin.** Segmentation of degenerating primary cortical axons treated with 200 µM hemin for 24 hours and recorded by time-lapse microscopy in a 30-minutes interval and shown in 7 frames/second. Scale bar: 100 µm.
