## Supplementary figures and images for "Deep learning to decipher the progression and morphology of axonal degeneration"

### Video S1

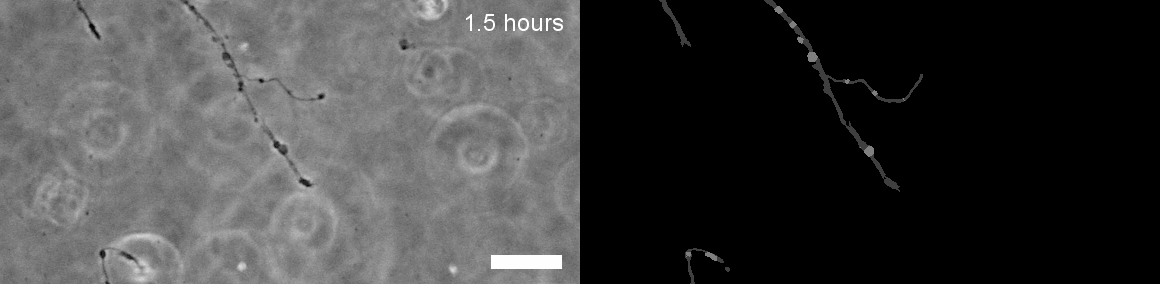

### Video S2

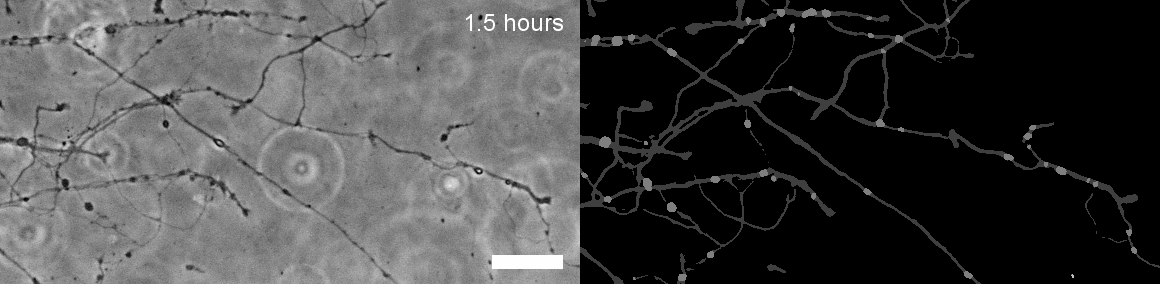

### Video S3

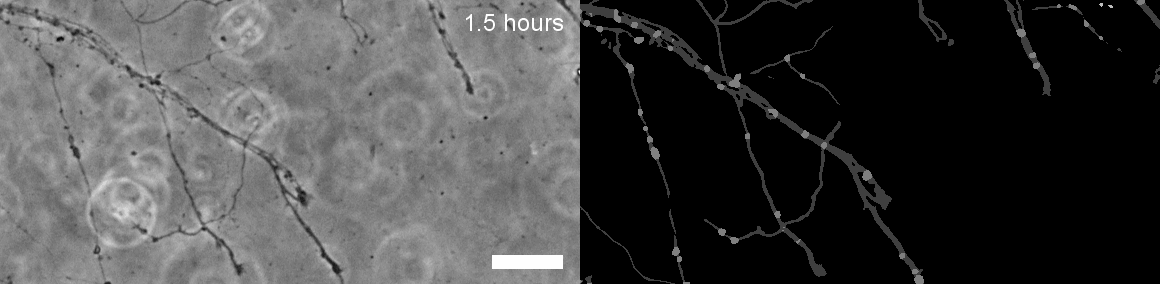

### Video S4

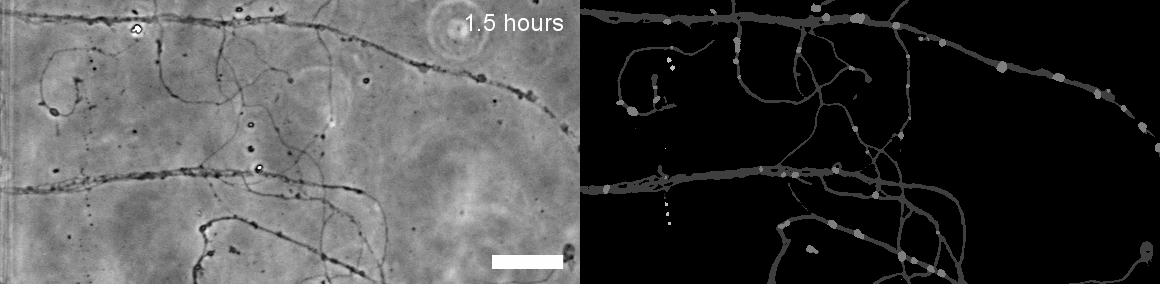

### Video S5

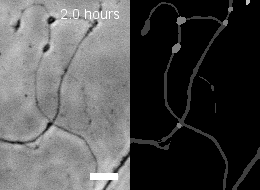

### Video S6

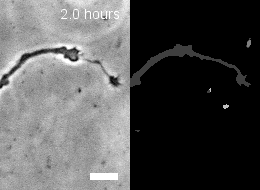

### Video S7

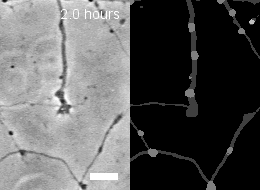

### Video S8

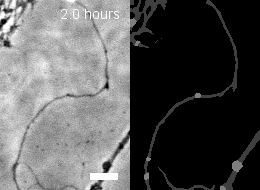

### Video S9

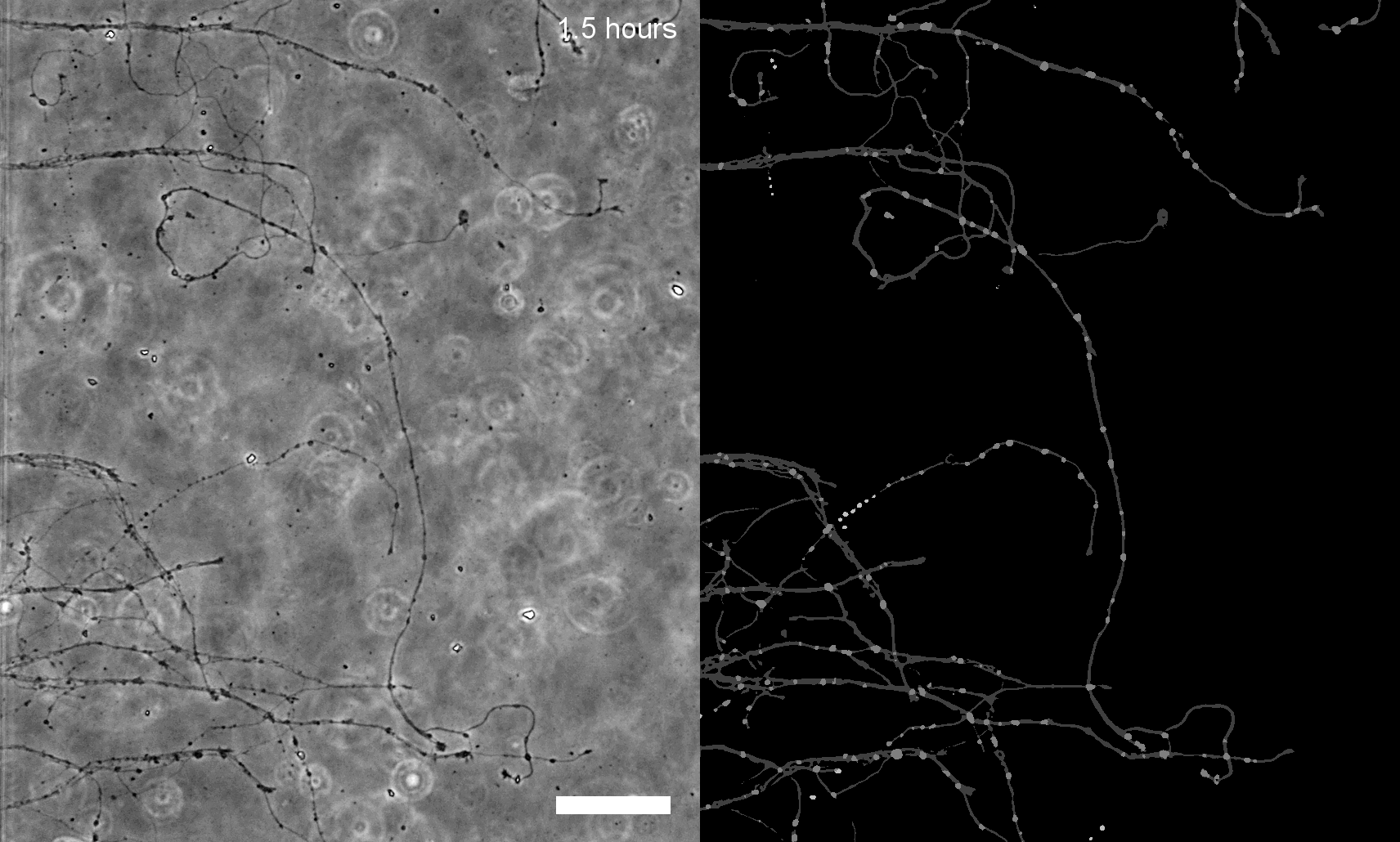
